# TFAP2 paralogs regulate midfacial development in part through a conserved *ALX* genetic pathway

**DOI:** 10.1101/2023.06.16.545376

**Authors:** Timothy T Nguyen, Jennyfer M Mitchell, Michaela D Kiel, Kenneth L Jones, Trevor J Williams, James T Nichols, Eric Van Otterloo

## Abstract

Cranial neural crest development is governed by positional gene regulatory networks (GRNs). Fine-tuning of the GRN components underly facial shape variation, yet how those in the midface are connected and activated remain poorly understood. Here, we show that concerted inactivation of *Tfap2a* and *Tfap2b* in the murine neural crest even during the late migratory phase results in a midfacial cleft and skeletal abnormalities. Bulk and single-cell RNA-seq profiling reveal that loss of both *Tfap2* members dysregulated numerous midface GRN components involved in midface fusion, patterning, and differentiation. Notably, *Alx1/3/4* (*Alx*) transcript levels are reduced, while ChIP-seq analyses suggest TFAP2 directly and positively regulates *Alx* gene expression. *TFAP2* and *ALX* co-expression in midfacial neural crest cells of both mouse and zebrafish further implies conservation of this regulatory axis across vertebrates. Consistent with this notion, *tfap2a* mutant zebrafish present abnormal *alx3* expression patterns, and the two genes display a genetic interaction in this species. Together, these data demonstrate a critical role for TFAP2 in regulating vertebrate midfacial development in part through ALX transcription factor gene expression.

## INTRODUCTION

The evolution of paired jaws in parallel with the development of neural crest cells (NCCs) has led to considerable breadth of facial shapes, enabling gnathostome vertebrates to thrive and exploit varied ecological niches (Brandon et al., 2022; Martik and Bronner, 2021; Square et al., 2017; York and McCauley, 2020). The ability for cranial NCCs (CNCCs) to give rise to skeletal elements important for mastication and housing numerous sensory organs is driven by coordination of epigenetic priming and transcriptional responses to local signaling inputs during embryonic development (Clouthier et al., 2010; Kessler et al., 2023; Santagati and Rijli, 2003; Selleri and Rijli, 2023). Fine-tuning of these positional gene regulatory networks (GRNs) underscores facial shape variation; likewise, disruption of such important processes can lead to severe craniofacial birth defects including orofacial clefting. In humans, this phenotype usually presents as a lateral or bilateral cleft of the upper lip and primary palate which presents in ∼1 in 700 births (Mai et al., 2019; Tolarova and Cervenka, 1998). However, clefting can also occur at the midline of the medial- and upper-face region (i.e., midface) (Tessier, 1976), generating a bifid nose that is often part of the spectrum of pathologies such as Frontonasal Dysplasia (FND) (MIMs: 136760, 613451, 613456) (Vargel et al., 2021). Genes encoding the homeobox-containing ALX transcription factors (*ALX1/3/4*) are the prominent underpinning of FND when mutated and have also been linked with midfacial development in mouse and zebrafish (Beverdam et al., 2001; Iyyanar et al., 2022; Lakhwani et al., 2010; McGonnell et al., 2011; Mitchell et al., 2021; Qu et al., 1999; Yoon et al., 2022). Although *ALX* genes are clearly fundamental components of the GRNs regulating midfacial development, how they connect with other midface genes into regulatory nodes remains uncertain. Of further importance, midface-specific modules are postulated to drive facial shape variation in humans (Claes et al., 2018; Feng et al., 2021; Naqvi et al., 2022; Xiong et al., 2019). Therefore, uncovering and connecting these genes in the midface GRN remains central to understanding facial development, morphology, and pathology.

Encoded members of the *TFAP2* gene family have previously been linked with facial development, particularly as transcription factors operating in GRNs across multiple NCC lineages in several vertebrate species (de Croze et al., 2011; Dooley et al., 2019; Hovland et al., 2022; Kenny et al., 2022; Li and Cornell, 2007; Rothstein and Simoes-Costa, 2020; Schmidt et al., 2011; Seberg et al., 2017; Van Otterloo et al., 2010; Van Otterloo et al., 2012). TFAP2 paralogs share highly conserved amino acid sequences forming their DNA-binding and dimerization domains; moreover, their overlapping DNA-binding motifs (5’-GCCNNNGGC-3’) and capacity to form homo- or heterodimers suggest functional redundancy (Eckert et al., 2005; Williams et al., 1988; Williams and Tjian, 1991). In particular, TFAP2A and TFAP2B have been shown to regulate craniofacial development in numerous model organisms (Barrallo-Gimeno et al., 2004; Knight et al., 2003; Martino et al., 2016; Schorle et al., 1996; Zhang et al., 1996) and are also mutated in human syndromes with facial pathologies. Dominant mutations in *TFAP2A* and *TFAP2B* cause Branchio-Oculo-Facial syndrome (BOFS) (MIM: 113620) (Milunsky et al., 2008) and Char syndrome (MIM: 169100) (Satoda et al., 2000), respectively, conditions which include midface phenotypes such as hypertelorism, broad nasal tip, as well as an abnormal philtrum and nasal bridge. TFAP2 has also been implicated as a central determinant to facial evolution (Prescott et al., 2015) and, thus, facial morphology appears particularly sensitive to the total functional levels of TFAP2 (Laugsch et al., 2019; Naqvi et al., 2022). Supporting this notion, biochemical analyses suggest the BOFS and Char syndrome mutations interfere with the function of the remaining wild-type product or other TFAP2 paralogs (Li et al., 2013; Satoda et al., 2000). Moreover, simultaneous targeting of *TFAP2A* and *TFAP2B* in mice and zebrafish results in exacerbated craniofacial defects (Knight et al., 2005; Van Otterloo et al., 2018; Van Otterloo et al., 2022; Woodruff et al., 2021). Mechanistically, TFAP2 orchestrates branchial arch (BA) patterning through modulating homeobox transcription factor gene expression (Knight et al., 2004; Knight et al., 2003; Van Otterloo et al., 2018), thus raising the possibility that similar mechanisms operate in the midface.

Here, we coupled mouse and zebrafish genetics with next-generation sequencing to extend our analysis of TFAP2A and TFAP2B function in midfacial development and gene regulation. Our findings show that similar midfacial clefting phenotypes are observed when these two genes are lost either early or late in mouse NCC development, suggesting that the critical aspects of their function reside in patterning and differentiation rather than migration. Using genome-wide molecular analysis in mouse coupled with genetic approaches in zebrafish, we further link these TFAP2 genes with ALX family expression and function, placing them firmly in a GRN critical for midface development and evolution.

## RESULTS

### TFAP2A and TFAP2B cooperatively function in CNCCs during midface development

In our previous efforts to illuminate the combined role of *TFAP2A* and *TFAP2B* in CNCCs, we identified that simultaneous loss of these genes in mice caused major defects in the development of both midface and BA1-derived jaw structures (Fig. S1A-F) (Van Otterloo et al., 2018). Subsequent profiling focused on the disruption of GRNs patterning BA1, while mechanisms responsible for the midfacial cleft were not pursued in detail. These earlier studies also deployed a combination of null and floxed conditional alleles such that lack of *Tfap2* in NCCs was accompanied by their heterozygosity in the remaining embryo (Fig. 1A). Since the concerted action of TFAP2A and TFAP2B serves critical functions in the ectoderm during facial development (Van Otterloo et al., 2022; Woodruff et al., 2021), we wished to ascertain if the midfacial clefting was caused solely by loss of the two genes in NCCs.

**Fig. 1.**
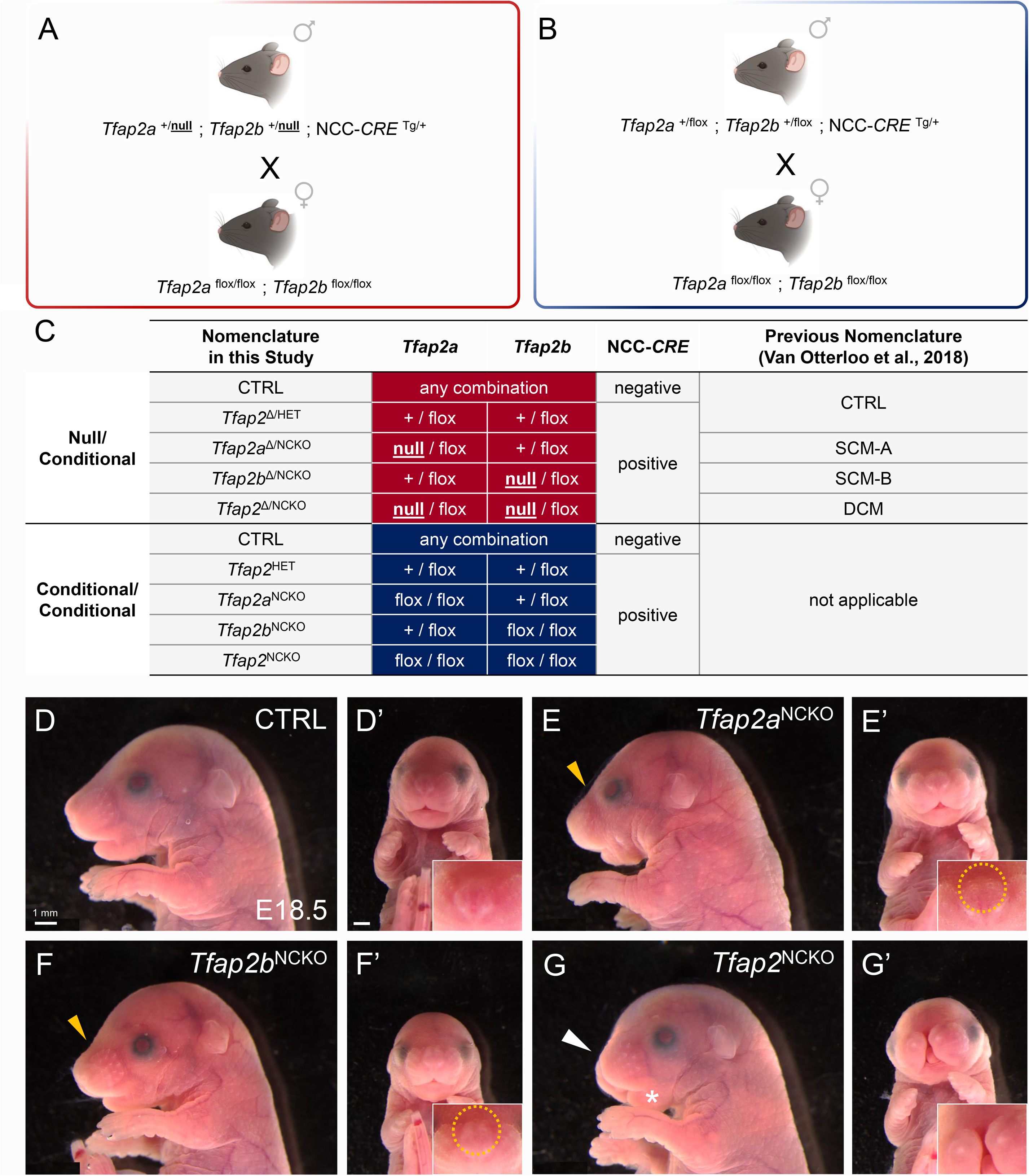
TFAP2A and TFAP2B cooperatively function in CNCCs during midfacial development. **(A, B)** Schematics depicting the mouse breeding scheme used for the null/conditional model (A), previously described (Van Otterloo, et al., 2018), and the conditional/conditional model (B) used in this study. Cartoons are adapted from BioRender. **(C)** Table summarizing a subset of the combination of alleles that can be obtained using the breeding scheme highlighted in panels A or B, along with current and previous shorthand nomenclature. Note, null/conditional genotypes are denoted with a “Δ” superscript in the current study. **(D-G)** Lateral (D-G) or frontal (D’-G’) views of conditional/conditional mouse embryos at embryonic day 18.5 (E18.5), with indicated genotypes. Gold arrowheads in panels E and F point to indented snouts. Insets in panels D’-G’ include higher magnification images of the snout, with misplaced vibrissae outlined in a gold dashed circle. White arrowhead in G indicates the shortened snout and the asterisk marks the short mandible. N = 5 per genotype.

To address this, we utilized only floxed conditional alleles in combination with NCC-specific *CRE* recombinase transgenes (Fig. 1B). Moving forward, embryos derived from the prior breeding scheme (Fig. 1A) are denoted as the null/conditional model (Fig. 1C, red), while embryos from the updated paradigm (Fig. 1B) are termed as the conditional/conditional model (Fig. 1C, blue). First using *Wnt1:CRE* (Danielian et al., 1998) in the updated paradigm, we conditionally deleted various allelic combinations of *Tfap2* genes in pre-migratory NCCs and examined gross craniofacial development (Fig. 1C-F). *CRE*-negative control mice at embryonic day 18.5 (E18.5) displayed appropriate midface formation that included laterally positioned vibrissae and an elongated snout (Fig. 1D). Littermates with NCC-specific loss of *Tfap2a* (*Tfap2a*^NCKO^, also *Tfap2b*-heterozygous) or *Tfap2b* (*Tfap2b*^NCKO^, also *Tfap2a*-heterozygous) displayed mild and largely overlapping midfacial anomalies such as an indented snout (Fig. 1E, F, gold arrowhead) with misplaced vibrissae on the dorsum (Fig. 1E’, F’, circled). By contrast, complete loss of *Tfap2a* and *Tfap2b* in NCCs (*Tfap2*^NCKO^) resulted in a fully penetrant midfacial cleft that was accompanied by a mildly shortened snout (Fig. 1G; white arrowhead). The misplaced vibrissae present by loss of a single paralog were not observed in *Tfap2*^NCKO^ embryos, but we surmised this was confounded by the facial cleft. Earlier timepoints (E11.5, E12.5, E15.5) revealed similar phenotypic trends, with the cleft becoming evident at E11.5 (Fig. S2) during midface fusion.

Jaw defects were also apparent in the various mutants by gross examination or following skeletal analysis (Fig. S3), resembling those observed in our prior studies (Van Otterloo et al., 2018). Intriguingly, the conditional/conditional embryos did not display a midline mandibular cleft, in contrast to previous findings in their *Tfap2*^Δ/NCKO^ counterparts (Fig. S1). Therefore, the current analysis indicates that the upper midface cleft and significant BA1 patterning defects can be attributed solely to loss of these two genes in NCCs, whereas mandibular clefting is uncovered by the reduced allelic dosage of *Tfap2* elsewhere— presumably the ectoderm.

In summary, anomalies like the midfacial cleft suggest that proper facial development relies on sufficient *Tfap2* dosage specifically in NCCs. As observed in other NCC stages and derivatives (Rothstein and Simoes-Costa, 2020; Schmidt et al., 2011; Seberg et al., 2017; Van Otterloo et al., 2018), these findings raise the hypothesis that TFAP2A and TFAP2B cooperatively function in NCC GRNs directing midfacial development.

### Fusion of the frontonasal prominences require TFAP2 function in post-migratory CNCCs

TFAP2 paralogs are postulated to be critical pioneer transcription factors residing throughout NCC development, marking genes poised to be activated once CNCCs have taken residency in the facial prominences (Fernandez Garcia et al., 2019; Rada-Iglesias et al., 2012; Rothstein and Simoes-Costa, 2020). We next asked whether the midfacial cleft resulted from the lack of TFAP2 activity throughout NCC development or after their migration. Previously assessed by β-galactosidase staining with the *r26r-lacZ* reporter (Soriano, 1999), E9.0 and E10.0 *Tfap2*^Δ/NCKO^ embryos revealed no distinct changes in CNCC migration into the facial complex (Van Otterloo et al., 2018), which comprise of the midface-associated frontonasal prominence (FNP) alongside the paired maxillary prominences (MxP) a mandibular prominences (MdP). Extending this analysis in conditional/conditional animals, we observed that the distribution and intensity of β-galactosidase staining in the E10.5 midface was similar between *Tfap2*^NCKO^, *Tfap2a*^NCKO^, *Tfap2b*^NCKO^, and control littermates heterozygous for *Tfap2* in NCCs (*Tfap2*^HET^) (Fig. S4A), with the caveat that the developing cleft between the FNP alters overall morphology in *Tfap2*^NCKO^ embryos (Fig. S4A, arrowheads).

Performing sectional immunofluorescence on E11.5 wild-type midface tissue, we found TFAP2A and TFAP2B protein expression in CNCCs occupying the medial and lateral domains of the FNP (Fig. 2A; Fig. S4B). Additional comparisons of published E11.5 transcriptomes showed persistent *Tfap2a* and *Tfap2b* transcript levels in FNP CNCCs in contrast to those in the MdP (Fig. 2B) (Hooper et al., 2020), Therefore, taking advantage of the *Sox10:CRE* transgene (Matsuoka et al., 2005) to inactivate *Tfap2* when CNCCs have begun reaching the FNP (∼E9.0) (Hari et al., 2012; Jacques-Fricke et al., 2012), we next sought to determine the critical periods by which TFAP2 is needed for FNP fusion. Micro-computed tomography (µCT) analysis at E12.5 showed that, indeed, similar to *Wnt1:CRE* animals (Fig. 2C), *Sox10:CRE-Tfap2*^NCKO^ embryos presented a midface cleft not observed in controls (Fig. 2D). Quantification of the linear distance between medial FNP tips were statistically significant between controls and their respective mutant backgrounds (Fig. 2E), with *Sox10:CRE*-*Tfap2*^NCKO^ more mildly affected at this stage, while ventral µCT sections revealed that both mutants shared abnormal nasal pit morphology (Fig. 2F, G). Lastly, examining the gross morphology of E18.5 *Sox10:CRE* embryos of the various *Tfap2* allelic combinations presented similar phenotypic trends as their *Wnt1:CRE* counterparts (Fig. S5).

**Fig. 2.**
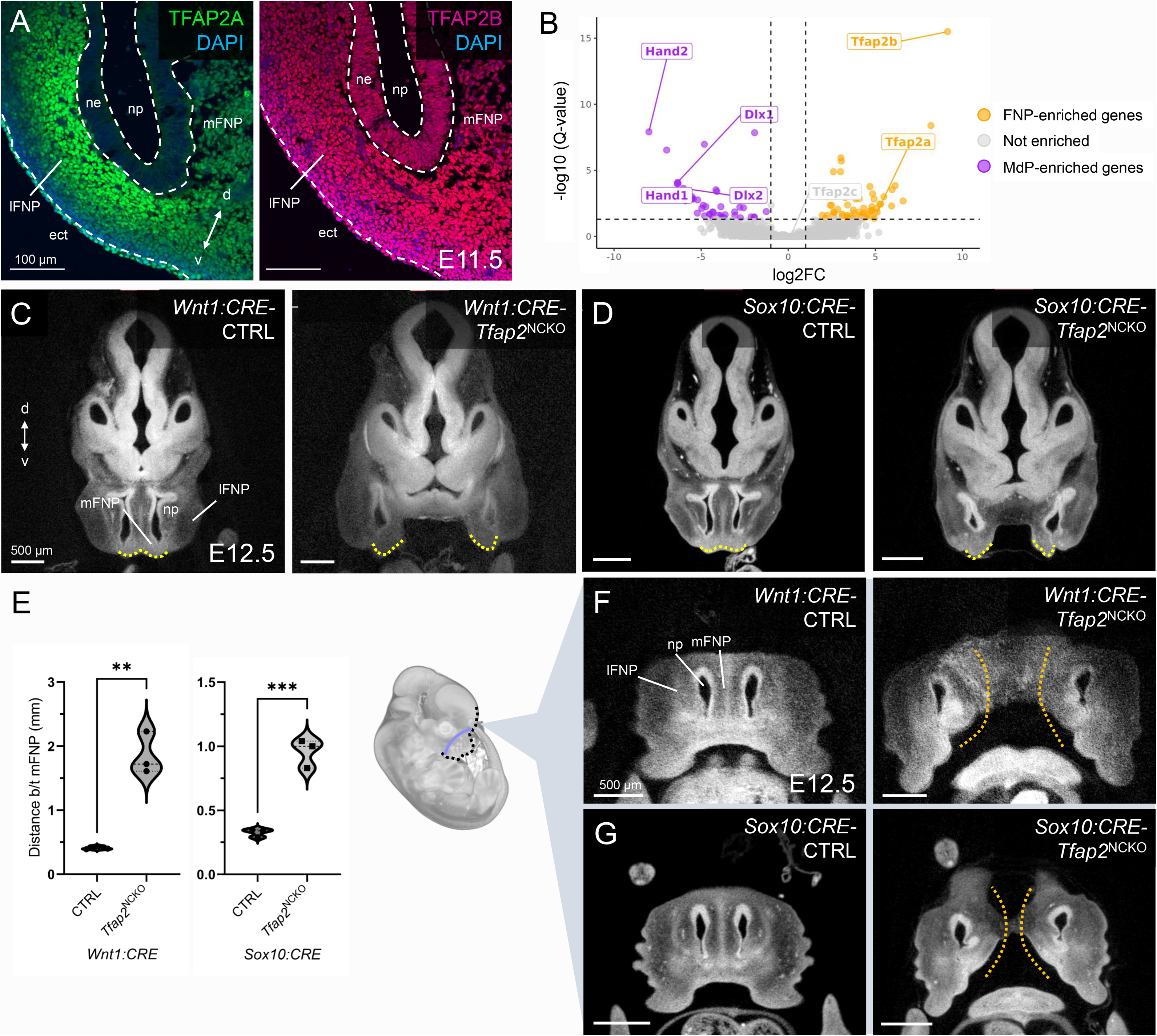
TFAP2A and TFAP2B function during post-migratory CNCCs to facilitate midface fusion. (**A**) Immunofluorescent analysis of TFAP2A (left, green) and TFAP2B (right, red) on a horizontal tissue section through an E11.5 wild type embryonic midface. White dashed lines demarcate ectoderm-mesenchyme boundaries. (**B**) Volcano plot displaying differentially expressed genes between E11.5 CNCCs occupying the mandibular prominence (MdP, violet) and those occupying the frontonasal prominence (FNP, orange). (**C, D**) µCT horizontal sections of E12.5 control-*Tfap2*^NCKO^ littermates in either the *Wnt1:CRE* (C) or *Sox10:CRE* (D) models. Yellow dashed lines outline the medial domains of the frontonasal prominence (mFNP). (**E**) Quantification of the linear distance between the tips of the mFNP domains in control and *Tfap2*^NCKO^ embryos in both mouse models. N = 3 per genotype. Student’s t-test, ** p < 0.01, *** p > 0.001. (**F, G**) µCT frontal sections through the snout. Gold dashed lines highlight the midfacial cleft. Additional abbreviations: d, dorsal; ect, ectoderm; lFNP, lateral FNP; ne, nasal epithelium; np, nasal pit; v, ventral.

Thus, while TFAP2 persists throughout CNCC developmental progression, these data provide evidence that the midface cleft does not arise from impaired TFAP2 activity during migration. Rather, retention of TFAP2A and TFAP2B expression levels in post-migratory CNCCs, alongside the *Sox10:CRE* studies, refine the temporal requirement for both paralogs in midface fusion.

### Appropriate midfacial bone and cartilage formation depends on *Tfap2a* and *Tfap2b* gene dosage

We next sought to leverage our transgenic models to better understand the developmental timing of TFAP2 during midfacial bone and cartilage formation. Accordingly, we compared E18.5 skeletal preparations of *CRE*-negative control, *Tfap2a*^NCKO^, *Tfap2b*^NCKO^, and *Tfap2*^NCKO^ backgrounds from both conditional/conditional paradigms. Detailing the midfacial skeleton first in *Sox10:CRE* animals, controls showed appropriate ossification of frontal and nasal bones meeting at the midline (Fig. 3A). Underneath the cranium lies the primary palate formed by the premaxilla bones, while the anterior portions of the vomer bones meet at the midline (Fig. 3A’). The nasal capsules are joined, connected to a compact cartilaginous template for the nasal (i.e., ethmoid) labyrinth and nasal septum that sits adjacent to the anterior presphenoid (Fig. 3A’). *Tfap2a*^NCKO^ and *Tfap2b*^NCKO^ backgrounds exhibited small ectopic cartilage islands forming on the calvaria, mildly increased gaps between the frontal bones (Fig. 3B, C). Comparing *Tfap2a*^NCKO^ and *Tfap2b*^NCKO^ mutants, the former exhibited gaps in the nasal bones and mild split in the vomer bones (Fig. 3B, B’, white arrowhead, C, C’). By contrast, *Tfap2*^NCKO^ embryos lacked nasal bones, exposing the underlying split nasal capsules (Fig. 3D, asterisk). Large cartilaginous ectopias were also found in between the hypoplastic frontal bones that, in some individuals, appeared to stem from the nasal septum (Fig. 3D, gold arrowheads; Fig. S6A, A’). Interestingly, in one individual these ectopias were replaced by severe frontal bone fractures (Fig. S6A”). Underneath the calvaria, *Tfap2*^NCKO^ mutants presented clefting of the palatal processes of the premaxilla, increased separation between the split vomer bones, less compacted nasal labyrinths, a thickened nasal septum, and malformed presphenoid bone (Fig. 3D’; Fig. S6B-B”, dashed lines). Although jaw and cranial base defects were not closely examined, it is worth mentioning that major BA1 defects were in line with those observed in both *Wnt1:CRE* animals (Fig. S7) (Van Otterloo et al., 2018). Intriguingly, only one *Tfap2*^NCKO^ mutant presented fusion of the paired jaws (i.e., syngnathia) (Fig. S7E), a defect with higher prevalence in *Wnt1:CRE-Tfap2*^NCKO^ and *Tfap2*^Δ/NCKO^ embryos (Fig. S3) (Van Otterloo et al., 2018). Overall, the more common phenotype was fusion between maxillary and jugal bones (Fig. S7F).

**Fig. 3.**
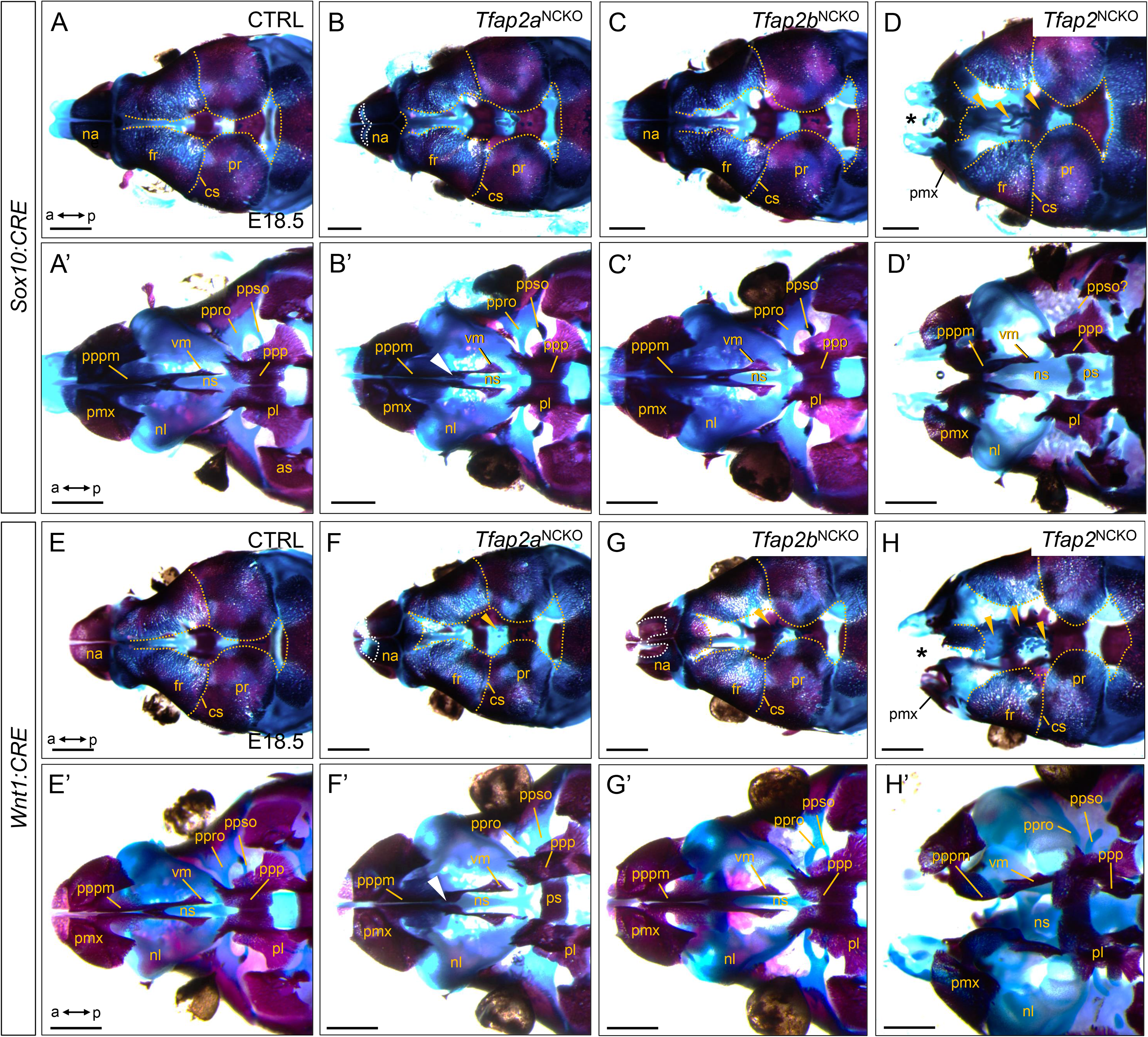
Formation of midfacial bone and cartilage structures depends on Tfap2a and Tfap2b gene dosage in CNCCs. (**A-H**) Concurrent Alizarin Red/Alcian Blue stained skeletal preparations of E18.5 embryos, with indicated genotypes, from *Sox10:CRE* (**A-D**) or *Wnt1:CRE* (**E-H**) based breeding schemes. Anterior (a) is to the left. Bone is stained magenta while cartilage is stained blue. However, note that cartilage staining towards the anterior end is variable that precluded thorough examination of the nasal capsules. Top-down, dorsal, (**A-H**) or bottom-up, ventral, (**A’-H’**) views of the midfacial skeleton, with the maxillary bone removed. Gold dashed lines (**A-H**) outline the peripheral edges of the calvarial bones. White dashed lines (**B, F, G**) outline nasal bone gaps. Gold arrowheads (**D, F-H**) point to cartilaginous ectopias sitting adjacent to the frontal bones. White arrowheads (**B’, F’**) point to a gap between the anterior portion of the vomer bones. Asterisks (**D, H**) highlight missing nasal bones. N = 5 per genotype. Scale bar = 1 mm. Additional abbreviations: cs, coronal suture; fr, frontal bone; na, nasal bone; ns, nasal septum; nl, nasal/ethmoid labyrinth; p, posterior; pl, palatine bone; pmx, premaxilla; ppp, palatal process of the palatine; pppm, palatal process of the premaxilla; ppro, pila preoptica; ppso, pila postoptica; pr, parietal bone; ps, presphenoid; vm, vomer bone.

Next examining the midface elements in *Wnt1:CRE* animals, both *Tfap2a*^NCKO^ and *Tfap2b*^NCKO^ single mutants presented largely overlapping phenotypes as their *Sox10:CRE* equivalents, though with slight differences (Fig. 3E-G, E’-G’). For example, compared to controls (Fig. E, E’), both single conditional mutants share gapped nasal bones and mishappened cartilage scaffolds connecting the frontal bone and anterior cranial base (pila pre/postoptica) (Fig. 3F, G, F’, G’). Strikingly, *Wnt1:CRE-Tfap2*^NCKO^ embryos presented nearly identical phenotypes like those observed with their *Sox10:CRE* counterparts, including a complete lack of nasal bones, smaller frontal bones, cranial ectopias, and overt separation between the premaxillary palate and vomer bones (Fig. 3H, H’). Moreover, additional preparation of E15.5 and E18.5 *Tfap2*^Δ/NCKO^ skeletons displayed similar midfacial hypoplasia and ectopia phenotypes (Fig. S8). It is unclear whether subtle phenotypic distinctions between *Sox10:CRE* and *Wnt1:CRE* mice are confounded by differences in *CRE* spatial expression patterns (Debbache et al., 2018). Nevertheless, overlaps in major defects between animal models further emphasize critical dependence on *Tfap2a* and *Tfap2b* gene dosage during post-migratory CNCC events such as skeletal differentiation.

### Combined loss of *Tfap2a* and *Tfap2b* dysregulates a post-migratory midfacial neural crest GRN that includes Frontonasal Dysplasia-related *Alx1/3/4* genes

Our mouse genetic analyses motivated the hypothesis that TFAP2A and TFAP2B acted upstream at key post-migratory GRN modules associated with midface mesenchyme patterning and differentiation. Herein, we undertook transcriptomic profiling of controls and mutants during critical timepoints for midface fusion: RNA-seq of E10.5 bulk FNP/MxP tissue and scRNA-seq of E11.5 CNCCs, respectively. With respect to the first approach, comparison between *CRE*-negative control and *Tfap2*^Δ/NCKO^ mutants uncovered 149 genes to be dysregulated between groups (86 down-regulated, 63 up-regulated) (Fig. 4A; Table S1). Ontology, enrichment, and pathway analysis via the Enrichr pipeline (Kuleshov et al., 2016) revealed an overrepresentation of midfacial pathology categories associated with down-regulated genes (Fig. 4B). Similar to *Tfap2*^Δ/NCKO^ profiles in the lower jaw (Van Otterloo et al., 2018), many of the terms contained homeobox transcription factor genes including calvaria development-related *Msx1* (Han et al., 2007; Ishii et al., 2005; Roybal et al., 2010) as well as *Pax3* and *Pax7* (Gu et al., 2022; Zalc et al., 2015). Notably, terms like *midline defect of the nose* (p = 2.00e-05, adj. p < 0.05) as well as *Frontonasal Dysplasia* (6.33e-06, adj. p < 0.05) contained *Alx1*, *Alx3*, *Alx4* (Fig. 4A, B), whose combinatorial loss lead to a midface cleft strikingly similar to those in *Tfap2*^NCKO^ and *Tfap2*^Δ/NCKO^ (Fig. S2; Table S1) (Beverdam et al., 2001; Gu et al., 2022; Iyyanar et al., 2022; Lakhwani et al., 2010; Qu et al., 1999). Also reduced were *Crabp1* and *Aldh1b1*, encoding regulators of a retinoic acid pathway known to regulate midface fusion (Fig. 4A; Table S1) (Gao et al., 2021; Kennedy and Dickinson, 2012; Lohnes et al., 1994; Williams and Bohnsack, 2019; Wu et al., 2022). Unexpectedly, we noted increased expression of genes encoding collagens (e.g., *Col14a1*, *Col3a1*), periostin (*Postn*), osteoglycin (*Ogn*), and Fibronectin Leucine Rich Transmembrane Protein 2 (*Flrt2*) (Fig. 4A; Table S1), components involved in skeletal formation (Bhatt et al., 2013; Dash and Trainor, 2020) and suggest premature activation of this process. Thus, by E10.5, transcriptional changes are already emerging such as those involved in mammalian midfacial anomalies.

**Fig. 4.**
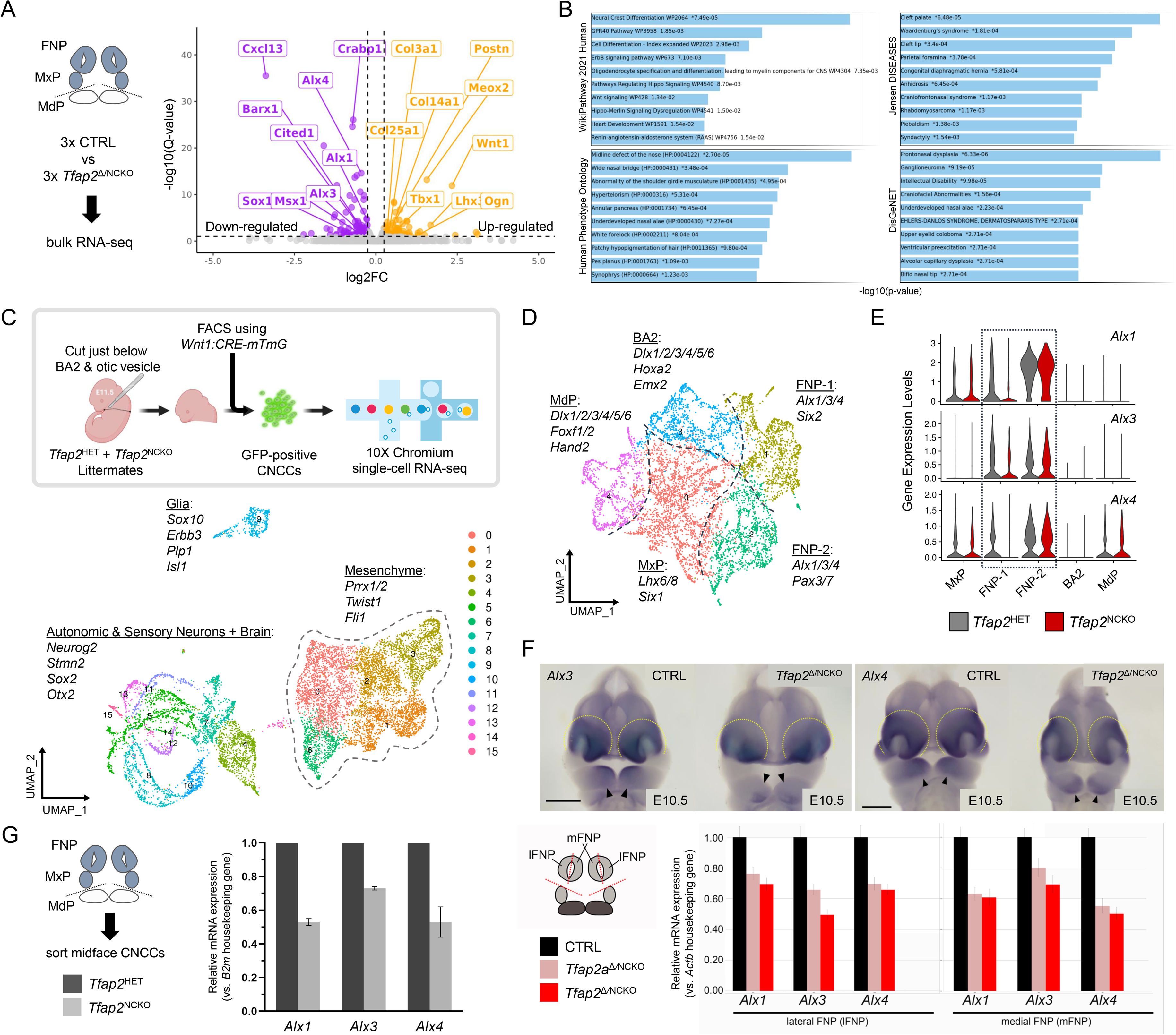
Transcriptomic analyses reveal NCC-specific loss of Tfap2 compromises midface Alx1/3/4 gene expression. (**A**) Bulk RNA-seq volcano plot analysis of frontonasal and maxillary prominence tissue (FNP, MxP) isolated from E10.5 control and *Tfap2*^Δ/NCKO^ littermates (cartoon workflow on the left). Down-regulated genes are visualized in purple while up-regulated genes are gold. (**B**) Enrichr (Kuleshov et al., 2016) terms based on the downregulated gene-set from the bulk RNA-seq analysis. (**C**) Overview of the scRNA-seq experiment performed on E11.5 *Tfap2*^HET^ and *Tfap2*^NCKO^ littermates. The top box displays the workflow of sorting GFP-positive head cranial neural crest cells (CNCCs) from branchial arch 2 (BA2) and upward. Cartoon partly derived from BioRender. Displayed at the bottom is the resulting Uniform Manifold Approximation and Projection (UMAP) plot of the three major groupings and genes enriched in each. Outlined is the mesenchyme population that is further subset and re-clustered (**D**) using MAGIC imputation (van Dijk et al., 2018). Dashed lines on the mesenchyme UMAP demarcate the identified positional identity clusters, with their respective gene signatures, of the head CNCCs. These include clusters for the FNP (two generated), MxP, mandibular prominence (MdP), and BA2. (**E**) Violin plots of *Alx1/3/4* gene expression between conditions and divided based on clusters. Boxed are the FNP clusters. (**F**) Whole mount *in situ* hyberization for *Alx3/4* between E10.5 controls and *Tfap2*^Δ/NCKO^ mutants, viewed in a frontal fashion. Scale bar = 500 µm. Dashed lines outline the FNP, while arrowheads point to the dysregulated *Alx3/4* staining in the MdP that was previously characterized (Van Otterloo et al., 2018). (**G**) Real-time PCR analysis for *Alx1/3/4* gene expression in the conditional/conditional model (left) and null/conditional model (right), with indicated genotypes and timepoints. Note that in the conditional/conditional model used sorted CNCCs from the FNP/MxP. Meanwhile, the null/conditional model examined expression profiles in bulk tissue isolated from individual medial and lateral domains of the FNP (mFNP, lFNP).

Secondarily conducting scRNA-seq of CNCCs sorted from the heads of E11.5 *Tfap2*^HET^ and *Tfap2*^NCKO^ littermates (Fig. 4C), we next examined the changing GRN architecture amidst midface fusion and the extent to which transcriptomic identities were compromised. Integration and Uniform Manifold Approximation and Projection (UMAP) plotting of combined control and mutant genotypes by Seurat (Hao et al., 2021) generated 16 clusters that divided into three major cell populations (Fig. 4C). Concordant with *Wnt1:CRE* cell labelling and other scRNA-seq studies (Jacques-Fricke et al., 2012; Soldatov et al., 2019), these cells clustered into groups for glia (e.g., *Sox10*, *Erbb3*, *Plp1*, *Isl1*; cluster 9), neurons and components of the brain (e.g., *Neurog2*, *Stmn2*, *Sox2*, *Otx2*; clusters 4, 5, 7, 8, 10-13, 15), and the mesenchyme (e.g., *Prrx1/2*, *Twist1, Fli1*; clusters 0-3, 6) (Fig. 4C, Fig. S9). While total cell numbers between samples were not drastically altered, mutant quantities for glial and mesenchymal populations were reduced (∼7-8% each) while neuronal populations increased (∼12%) compared to controls (Fig. S10A). Gene expression-based cell cycle scoring did not detect noticeable changes in cell cycle progression of these populations (Fig. S10B), indicating cellular profiles between genotypes are not due to altered proliferation states. While it is possible that TFAP2 participates in biasing CNCCs to a particular cell fate decision, we reasoned their loss alone is not sufficient to drive drastic fate switches akin to previously reported factors (Cox et al., 2012; Fan et al., 2021; Scerbo and Monsoro-Burq, 2020; Simoes-Costa and Bronner, 2016; Soldatov et al., 2019). Supporting this notion is no obvious expansion in neurons, previously tested by whole-mount anti-Neurofilament immunostaining (Van Otterloo et al., 2018).

To extend our bulk RNA-seq analyses, we implemented two scRNA-seq workflows in parallel for the mesenchyme group (Fig. 4C): a “pseudobulk” approach for differential gene expression analyses and MAGIC imputation (van Dijk et al., 2018) to annotate major positional identities. The former identified 383 dysregulated genes (124 down-regulated, 259 up-regulated; log fold-change threshold = 0.1, adj. p < 0.05), that contained overlaps with RNA profiles in the E10.5 midface (Table S1). For example, downregulated genes yielded several Enrichr terms associated with transcription factor gene regulation and craniofacial features associated with midface molecules (e.g., *Msx1, Pax3, Pax7, Alx1, Alx4*) (Fig. S11A, Table S1). Although *Alx3* did not appear in our pseudobulk, we suspect this is partly due to it being the lowest expressing paralog in the whole mesenchyme alongside the limited sequencing depth. Intriguingly, the upregulated GRN modules yielded members of the *Fox* transcription factor family (e.g., *Foxd1*, *Foxf2*, *Foxp1*, *Foxp2*), downstream targets of a Hedgehog signaling pathway often associated with midline facial disorders when precociously activated (Fig. S11B, Table S1) (Brugmann et al., 2010; Jeong et al., 2004; Xu et al., 2022; Xu et al., 2018). Further, we continued to observe several extracellular matrix terms resulting from the multiplicity of elevated collagen genes (e.g., *Col1a1*, *Col2a1*, *Col3a1*, *Col4a1*, *Col6a1*, *Col6a2*) (Fig. S11B, Table S1).

Confirming the positional specificity of the disrupted GRNs, we mapped expression levels of select genes on the computationally re-clustered mesenchyme UMAP, which comprises the presumptive FNP (FNP-1/2, clusters 1, 2; *Tfap2a/b*+, *Alx1/3/4*+, *Six2*+), MxP (cluster 0; *Six1*+, *Lhx6/8*+), MdP (cluster 0, 4; *Dlx1/2/3/4/5/6*+, *Hand2*+), and branchial arch 2 (BA2, cluster 3; *Hoxa2*+) (Fig. 4D; Fig. S12; Fig. S13). Consistent with previously reported lower face hypoplasia phenotypes (Van Otterloo et al., 2018), mutant cells were cumulatively underrepresented across MxP, MdP, and BA2 clusters (Fig. S14A, A’), accompanying the decreased expression of various *Dlx*, *Hand2*, and *Gsc* transcripts (Table S1; Fig. S14B). Lastly, of the two FNP clusters, mutant cells increased in FNP-1 (58.4% mutant/41.6% control) while decreasing in FNP-2 (26.8% mutant/73.2% control) (Fig. S14A, A’). These changes were coupled with reduced levels of *Alx1/3/4* (Fig. 4E), *Msx1*, and *Pax3/7*, as well as upregulation of collagen and *Fox* genes (Fig. S14C). Orthogonal approaches like whole mount *in situ* hybridization (Fig. 4F) and real-time PCR (Fig. 4G) analyses validated that, compared to their respective controls, both *Tfap2*^NCKO^ and *Tfap2*^Δ/NCKO^ mutants exhibited reduced midfacial *Alx* gene expression.

In total, profiling the loss of *Tfap2* in CNCCs by transcriptomic and target gene expression profiling suggests these transcription factors play crucial roles in regulating genes in the various positional programs. Importantly, these data support the hypothesis that TFAP2 paralogs function upstream of critical GRN nodes involved in midface fusion and skeletal formation, which include the Frontonasal Dysplasia-associated *Alx1/3/4* genes.

### TFAP2 plays a direct role in the midface GRN in part by occupying *Alx1/3/4* regulatory elements

Our genome wide transcriptomic analyses identified important genes that were dysregulated by loss of *Tfap2* in the midface CNCCs, but whether these were direct or indirect targets of these two transcription factors remained unclear. Therefore, we conducted ChIP-seq in E11.5 whole facial prominence tissue using an antibody that recognizes TFAP2A/B/C (Fig. 5A, data not shown). We identified 13,778 enriched regions (i.e., TFAP2 ‘peaks’) relative to input (Fig. 5B, Table S2), while antibody specificity was reflected by the TFAP2 consensus binding motif being the most significantly enriched (p = 1.00e-154) (Fig. 5C). Further analysis of TFAP2 binding coordinates revealed approximately 35% of them being in proximal promoters, 23% intronic, 16% intergenic, 9% in the 5’ UTR, while remaining peaks were found in additional regions of the genome (Fig. 5C). Since we used whole facial prominence tissue for these studies, these peaks presumably reflect unique or shared targets within the FNP, MxP, MdP, as well as those present in other tissues such as ectoderm and neuronal tissue. To focus on genes relevant to the midface, we cross-referenced these peak data with gene expression data for the E11.5 FNP (Hooper et al., 2020). Of all genes with detectable midface expression (e.g., E11.5 FNP mesenchyme FPKM > 0, N = 15,842), the genes associated with at least one TFAP2 peak (N = 8,754) were expressed in the midface at a significantly higher level than midface genes (N = 7,088) without a TFAP2 associated peak (p = 2.60e-256) (Fig. 5D, Table S2). Further, despite the unique features of different datasets (e.g., E10.5 vs E11.5, NCC-only vs whole face, etc.), ∼66% (76 of 115) of the genes dysregulated in our overlapping bulk and scRNA-seq dataset, including both upper- and lower-face specific, were associated with a TFAP2 peak (Table S2).

**Fig. 5.**
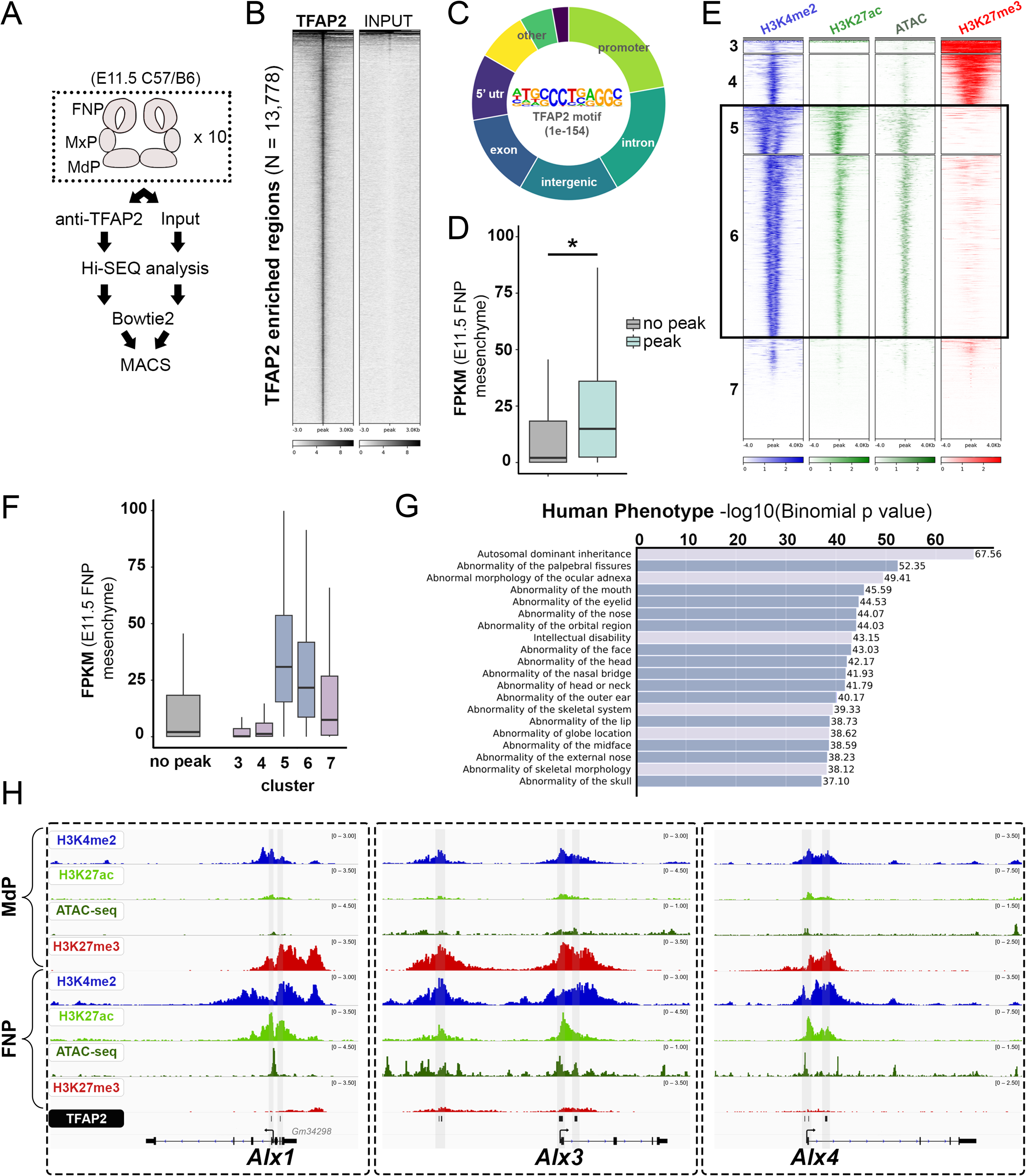
ChIP-seq profiling identifies TFAP2 paralogs directly bind Alx1/3/4 regulatory elements. (**A**) Schematic depicting workflow for anti-TFAP2 chromatin immunoprecipitation followed by sequencing (ChIP-seq) of wild-type E11.5 C57BL/6 facial prominence tissues. (**B**) Density heatmap displaying read-depth at the 13,778 TFAP2 ‘peaks’ (y-axis) detected by anti-TFAP2 ChIP-seq, relative to non-immunoprecipitated input. (**C**) A hollow pie chart summarizing the distribution of TFAP2 ‘peaks’ throughout the genome, relative to key features (e.g., promoters, introns, etc.). The TFAP2 motif, along with the significance of its enrichment in all TFAP2 ‘peaks’, is displayed in the center. (**D**) Box-and-whisker graph plotting the FPKM values of 16,573 genes with frontonasal prominence (FNP) mesenchymal expression at E11.5 (Hooper et al., 2020), with or without an associated TFAP2 peak (p-value = 2.60e-256). (**E**) Density heatmaps for anti-H3K4me2 (blue), anti-H3K27ac (green), ATAC-seq (forest green), and anti-H3K27me3 (red) profiles of E10.5 FNP cranial neural crest cells (CNCCs) (Minoux et al., 2017) at the TFAP2-positive coordinates. These regions are divided by k-means clustering. Boxed are clusters 5 and 6, whose coordinates are correlated with high H3K4me2, high H3K27ac, high ATAC (i.e., chromatin accessibility), and low H3K27me3 signals. (**F**) Box-and-whisker graphs, as in panel D, but further partitioning the ‘peak’ group based on k-means clusters from panel E. (**G**) GREAT (McLean et al., 2010) pathway analysis of genes associated with cluster 5 TFAP2 peaks. Significant “Human Phenotype” terms are listed, with the most significant on the top. Navy shaded boxes are associated with craniofacial-specific terms. (**H**) IGV browser views of the *Alx1*, *Alx3*, and *Alx4* loci overlayed with the epigenome signatures (blue, H3K4me2; green, H3K27ac; forest green, ATAC; red, H3K27me3) of E10.5 CNCCs residing in the mandibular prominence (MdP, top 4 tracks) and FNP (tracks 5-8), as well as the cluster 5-assigned TFAP2 peaks (bottom, gray highlights).

We next assessed chromatin states at these ∼13,800 genomic regions with the prediction that a subset of these coordinates would be found within FNP expressed, CNCC-associated, accessible, and ‘active’ regions of the genome. Using k-means clustering along with previously published histone ChIP-seq and chromatin accessibility datasets generated from wild-type E10.5 FNP CNCCs (Minoux et al., 2017), 5 major clusters (clusters 3, 4, 5, 6, 7; cluster 1 and 2 had < 6 peaks) were identified (Fig. 5E, Table S2). Of these five clusters, over half of TFAP2 bound regions (N = 7,995; 58% of all peaks) clustered into 2 major groups (clusters 5, 6) (Fig. 5E, Table S2). Both clusters contained substantial levels of the active H3K4me2 and H3K27ac, accessible chromatin, and marked reduction in the repressive H3K27me3 mark (Fig. 5E). In contrast to clusters 5 and 6, clusters 3 and 4 displayed substantially reduced levels for open chromatin and active H3K27ac accompanied by expanded H3K27me3 signal (i.e., repressed regions in E10.5 FNP CNCCs) (Fig. 5E, Table S2). Consistent with these profiles, genes associated with clusters 5 and 6 were highly expressed in the FNP, relative to genes without a TFAP2 peak, or genes associated with a peak(s) found in other clusters (Fig. 5F, Table S2). Pathway analysis for cluster 5 (1,658 peaks) and cluster 6 (6,336 peaks) revealed terms associated with head and midfacial pathology (Fig. 5G). Notably, genes associated with cluster 3 and 4 were significantly enriched for skin (cluster 4), neuronal (clusters 3, 4), and cardiac (cluster 3) terms (Fig. S15), potentially reflecting TFAP2 binding in non-NCC tissues in which TFAP2 is expressed (e.g., ectoderm). Finally, cluster 7 displayed relatively little FNP associated histone and chromatin signatures (Fig. 5E, Table S2) and displayed the fewest associated terms (data not shown).

Given the significant down-regulation of all three *Alx* genes in *Tfap2*^NCKO^ and *Tfap2*^Δ/NCKO^ mutants (Fig. 4) and the midfacial cleft associated with their loss (Beverdam et al., 2001; Iyyanar et al., 2022; Lakhwani et al., 2010; Qu et al., 1999), we next asked whether the *Alx* loci were directly bound by TFAP2. Assessment of peak files revealed that all 3 *Alx* loci had enriched regions of TFAP2 binding (Fig. 5H); moreover, these regions were associated with cluster 5, concordant with active and open genomic regions at the site of TFAP2 binding (Fig. 5H). Consistent with the more limited expression of *Alx1, Alx3,* and *Alx4* in jaw domains at E11.5, these genomic regions were associated with reduced H3K4me2 and H3K27ac and increased H3K27me3 in CNCCs occupying the MxP and MdP (Fig. 5H, and not shown) (Minoux et al., 2017). Thus, transcription factor binding analysis suggests that TFAP2 paralogs occupy genomic regions harboring active histone enhancer marks and associated with midfacial patterning and differentiation genes, namely *Alx* genes.

### Zebrafish *tfap2a* regulates *alx3* expression and genetically interacts with *alx3* during midfacial development

While different vertebrates use different TFAP2 paralogs during CNCC development, a member commonly used is TFAP2A (Eckert et al., 2005; Meulemans and Bronner-Fraser, 2002). Like humans and mice, the zebrafish *tfap2a* paralog is a critical determinant for formation of the craniofacial structures (Neuhauss et al., 1996; Schilling et al., 1996). Mining our published CNCC scRNA-seq data (Mitchell et al., 2021; Stenzel et al., 2022), we observed *tfap2a* gene expression predominantly in the frontonasal cluster giving rise to the neurocranium (i.e., midface skeleton) (Fig. S16). Further co-expression analyses indicated that *tfap2a-*positive cells also express *alx3* (Fig. S16), suggesting that TFAP2A regulation of *ALX* gene expression is a conserved transcriptional axis in vertebrates. To test this, we examined *alx3* expression in wild-types and *tfap2a* mutants (Fig. 6A-D”) (Knight et al., 2003; Schilling et al., 1996). We analyzed zebrafish embryos using *in situ* hybridization at 48 hours post-fertilization, a stage resembling the mouse E10.5 stage when changes in *Alx* expression were identified in mouse *Tfap2a*^Δ/NCKO^ FNP tissue (Fig. 4G). In wild-type embryos, *alx3* expression is enriched in *fli1a:EGFP*-labelled CNCCs on the developing roof of the mouth (Fig. 6A-A”, C-C”), as well as those ventral and medial to the nasal placodes (Fig. 6A-A”). In *tfap2a* mutants, *alx3* expression patterns were altered, becoming reduced around the periphery of the nasal placodes, and bifurcated into two laterally expanded domains (Fig. 6B, D, E). Similar to prior zebrafish BA studies (Barrallo-Gimeno et al., 2004) and our mouse study (Fig. S4A), CNCC number and positioning appeared to be largely intact in the frontonasal region in *tfap2a* mutants (Fig. 6A’-D’, A”-D”). These data strongly suggest that the transcriptional circuit identified in post-migratory CNCCs is conserved between mice and zebrafish.

**Fig. 6.**
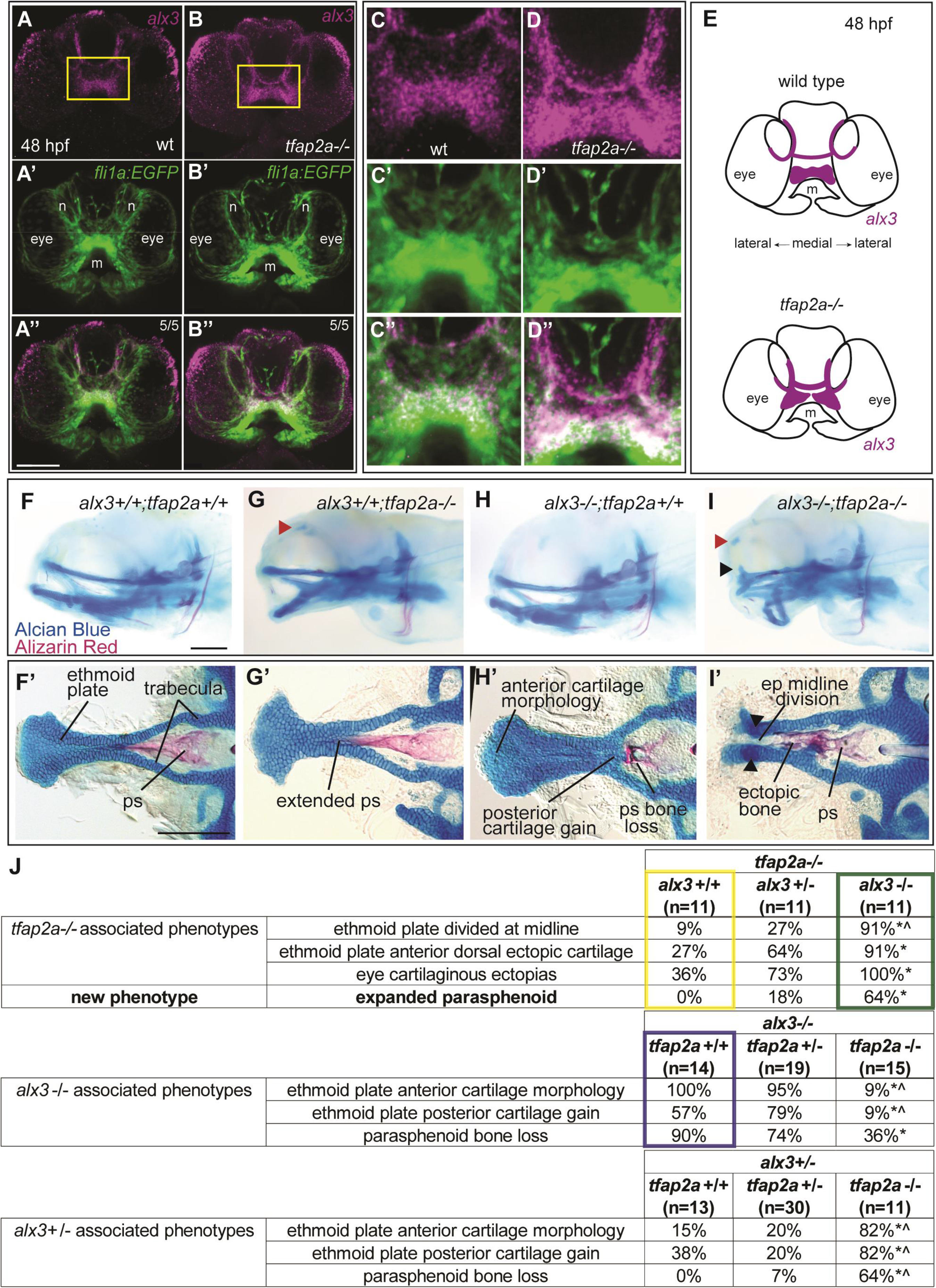
Gene expression and epistasis studies indicate zebrafish alx3 is genetically downstream of tfap2a. (**A-D**) Frontal views of *alx3 in situ* hybridization (violet) and *fli1a:EGFP*-labelled cranial neural crest cells (green) in a wild-type control (**A, C**) and a *tfap2a*-/- sibling (**B, D**) at 48 hours post-fertilization (hpf). Note, panels C and D are magnified views of the roof of the mouth, as boxed in panels A and B. (**E**) Schematic summarizing expression patterns observed, N = 5 per genotype. (**F-I**) Whole mount, lateral images of Alizarin Red/Alcian Blue stained skeletal preparations of 6-day-old zebrafish heads, with indicated genotypes. The anterior is to the left. Red arrowhead. (**F’-I’**) Neurocrania were dissected and flat mounted. Red arrowhead indicates ectopic cartilage near the eye socket. Black arrowhead marks dorsally projecting ectopic ethmoid plate cartilage. (**J**) Tables summarizing penetrance, samples, and statistics of the epistasis experiments as follow: top, the removal of wild-type *alx3* copies from *tfap2a* homozygous mutants; middle, the removal of wild-type *tfap2a* copies from *alx3* homozygous mutants; bottom, the removal of wild-type *tfap2a* copies from *alx3* heterozygotes. Boxed values correspond to the genotypes shown in panels G, H, and I. For example, in yellow are *tfap2a* mutant phenotypes and their penetrance (corresponding to G, G’); boxed in green those for when both *alx3* copies are removed in *tfap2a*-mutant backgrounds (corresponding to I, I’); lastly, boxed in purple are those for *alx3* mutants (H, H’). Asterisk indicates significant difference compared with the far-left column. Carets indicate significant difference compared with middle column by Fisher’s exact test. Scale bars = 100 µm. Abbreviations: ep, ethmoid plate; m, mouth; n, nares; ND, not determined; ps, parasphenoid.

Further testing the conservation hypothesis with the *tfap2a* and *alx3* mutant alleles (Mitchell et al., 2021), we sought to determine if the regulatory axis could be read out as genetic interactions (Fig. 6F-J). Several previous genetic studies indicate that viscerocranium structures derived from BA2-7 appear the most sensitive to *tfap2a* mutation, while the BA1 skeleton is less affected (Knight et al., 2003; Neuhauss et al., 1996; Schilling et al., 1996; Wang et al., 2011). In the anterior neurocranium, *tfap2a* mutants produced incompletely penetrant ethmoid plate loss with variable expressivity, ranging from complete absence to small discontinuations (Barrallo-Gimeno et al., 2004). In mutants with a divided ethmoid plate, the parasphenoid bone extended anteriorly, sometimes interfacing with the abnormal cartilage gap (Fig. 6G’) (Schilling et al., 1996). In this study, two incompletely penetrant *tfap2a* mutant-associated skeletal phenotypes were observed that were not previously described. Specifically, *tfap2a* mutant larvae exhibited partial penetrance for cartilaginous ectopias proximal to the eyes (Fig. 6G, red arrowhead) as well as ectopic cartilage rods projecting dorsally from the ethmoid plate (Fig. 6J). As *alx3* gene dosage was reduced in *tfap2a* mutants, the penetrance for all three phenotypes increased. We also discovered a new phenotype, never seen in single mutants, when copies of *alx3* were removed from *tfap2a* mutants (Fig. 6I, J). The parasphenoid is expanded in animals mutant for both *tfap2a* and *alx3.* It is interesting to note that a selection of these skeletal abnormalities parallel those observed in our mouse models. Specifically, the midfacial clefting and ectopic cartilage (Fig. 6I, I’) bear a striking resemblance to those found in *Tfap2*^NCKO^ and *Tfap2*^Δ/NCKO^ mutant embryos, suggesting that midline facial defects in both *TFAP2* mutant animal models are *ALX3*-dependent.

Performing the reciprocal genetic interaction experiment, we documented how *alx3* mutant-associated phenotypes were affected upon removing wild-type copies of *tfap2a. alx3* homozygous mutants present cellular morphology changes in the anterior ethmoid plate, an anteriorly truncated parasphenoid, and ectopic midline cartilage (Fig. 6J, middle table, boxed in purple) (Mitchell et al., 2021). In *alx3* mutant homozygotes with reduced *tfap2a* dosage, the ethmoid plate and parasphenoid were severely impacted (Fig. 6J, middle table), precluding us from scoring *tfap2a*-dependent changes. Therefore, we examined changes in *alx3-*associated phenotypes in *alx3* heterozygotes when wild-type copies of *tfap2a* were removed. Concordant with the first phenotypic scoring, we found that the removal of *tfap2a* copies from *alx3* heterozygotes increased penetrance of the *alx3* mutant phenotypes (Fig. 6J, bottom table). These findings demonstrate that *tfap2a* and *alx3* genetically interact during zebrafish midfacial development. Taken together with our observations in mice, these results raise a model that the transcription factors encoded by *TFAP2* and *ALX* genes function in a conserved, midface developmental pathway.

## DISCUSSION

### The role of TFAP2A and TFAP2B in cranial neural crest cells during midface development

Positional gene regulatory networks endow CNCCs their ability to form unique facial structures during embryogenesis. Despite the clear existence of GRN components directing midfacial development, shape, and pathology, our understanding of how they connect into transcription circuits is poorly understood. Our previous studies had established an essential role for NCC-specific *Tfap2a* and *Tfap2b* in facial development, and in particular, jaw patterning (Van Otterloo et al., 2018). While there are severe defects in both lower-face and midface regions, mechanisms underlying the latter were not investigated. In the current study, we show that TFAP2A and TFAP2B plays an essential role in CNCCs during midfacial development in part through activation of a conserved *ALX* genetic pathway. Deploying mouse models, we found that TFAP2-dependent midface fusion is largely driven by their regulatory control of post-migratory CNCCs. The combination of gene expression and transcription factor binding profiling suggest that TFAP2 paralogs act upstream of several GRN nodes critical for midface identity, midface fusion, and skeletal formation—notably through ALX transcription factor gene expression. Consistent with the notion that these regulatory mechanisms are conserved across jawed vertebrates, we found that zebrafish *tfap2a* is required for proper *alx3* expression patterns and that both genes interact in the midface skeleton in this species. Given the continuum of midface anomalies associated with disrupted *TFAP2A* (BOFS), *TFAP2B* (Char Syndrome), and *ALX* (FND) function in humans, these findings link features of previously disparate developmental disorders into a shared genetic hierarchy.

### TFAP2 and ALX paralogs in vertebrate midface morphology and evolution

Loss of *TFAP2* genes in CNCCs indicates a central role for the encoded proteins in defining positional neural crest GRNs, in part, by modulating homeobox transcription factor pathways. While TFAP2 is necessary for driving BA1 transcriptional programs—including *DLX* paralogs (Knight et al., 2003; Van Otterloo et al., 2018)—this study identified these factors execute a similar function in the midface. However, in this tissue, TFAP2 drives a unique repertoire of transcription factor genes, including *Pax3*, *Pax7*, *Msx1*, and of particular interest, *Alx1*, *Alx3*, and *Alx4*. Past studies using induced pluripotent stem-cell derived primate NCCs suggested TFAP2 and ALX proteins likely underlie species-specific facial variation (Prescott et al., 2015; Rada-Iglesias et al., 2012). More recent computational analyses of these datasets support the idea that *TFAP2* and *ALX* paralogs are linked to human facial morphology (Feng et al., 2021). Intraspecies comparisons further strengthen the role of both *TFAP2* [stickleback, (Erickson et al., 2018)] and *ALX* [Darwin’s finches, (Lamichhaney et al., 2015)] in facial evolution. Our *in vivo* ChIP-seq data suggests that the *Alx* loci (along with other lower-face and midface genes) are direct targets of TFAP2 binding. These findings are consistent with the study by Prescott *et al*., which postulated that human TFAP2A directly regulated genes hallmarking the various positional identities, including the *ALX* family (Prescott et al., 2015). Although additional studies are required for further detailing the relationship between TFAP2 and ALX factors, our *in vivo* analyses in mice and zebrafish recontextualize these *in vitro* studies, suggesting midface evolution is a direct output of gene regulation enacted by both transcription factor families.

### TFAP2 and ALX transcription factor pathways facilitating midface fusion

How might a TFAP2-driven ALX node in the midface GRN ultimately influence CNCC behaviors directing midface fusion? A combination of studies has provided some insight into the potential roles of ALX in this process. Most notably, ALX is a known regulator of skeletal differentiation across species. For example, in sea urchin, ALX directly targets biomineralization and extracellular matrix genes (Khor and Ettensohn, 2020). In line with these observations, our previous cellular and skeletal analyses in zebrafish implicate Alx3 as a critical determinant of midfacial chondrocyte differentiation timing (Mitchell et al., 2021). Similarly, in this study, we observed a significant increase in the gene expression of multiple collagens in *Tfap2*^NCKO^ mouse embryos (Fig. S11; Tables S1, S2)—whose deposition acts as a major readout of bone and cartilage formation (Dash and Trainor, 2020). Conceivably, aberrant activation of a differentiation GRN mediated via loss of TFAP2 could preclude appropriate midface fusion in an ALX-dependent manner.

There is precedent for ALX regulating additional aspects of post-migratory developmental programs. In the case of cell survival, *Alx1, Alx3,* and *Alx3/4* mutant mice exhibited elevated apoptosis in FNP CNCCs (Beverdam et al., 2001; Lakhwani et al., 2010; Zhao et al., 1996) and its periocular mesenchyme lineage (Iyyanar et al., 2022), while FND3 (*ALX1*) patient-derived NCC lines showed an increased propensity to cell death (Pini et al., 2020). Consistent with a role in patterning, *Alx1* and *Alx1/4* mutant mice exhibited a reduction of *Pax7* and an expansion of BA1-specific *Lhx6* and *Lhx8* into the FNP (Iyyanar et al., 2022). Interestingly, while a clear reduction in *Pax7* (and *Pax3*) expression was noted in *Tfap2*^Δ/NCKO^ and *Tfap2*^NCKO^ embryos, our expression profiling did not detect a concomitant expansion of *Lhx6* and *Lhx8*. The degree to which a reduction of *Alx* expression in *Tfap2* mutants contributes to the post-migratory CNCC phenotypes observed remains elusive and will require follow-up studies.

### Potential TFAP2 interplay with other midface positional factors

Our suite of animal and molecular studies suggests the TFAP2-ALX transcriptional circuit as a major node operating in the midfacial CNCCs. Yet, also evident was the disruption of additional GRN components, such as signaling pathways and other transcription factors. For example, loss of *Tfap2* resulted in decreased levels of various retinoic acid-related genes (Fig. 4A; Table S1, S2). This is of interest because loss of genes encoding components of the retinoic acid signaling pathway also results in a midline cleft (Gao et al., 2021; Kennedy and Dickinson, 2012; Lohnes et al., 1994; Wu et al., 2022), while the morphogen itself can modulate *TFAP2A* gene expression and activity *in vitro* (Luscher et al., 1989). Thus, TFAP2 may function in a regulatory feedback loop with this signaling cascade. Another pathway that is critical for midface development and that appeared to be disrupted upon loss of *Tfap2* included Sonic hedgehog signaling. While our gene expression profiling did not reveal significant changes in major Hedgehog signaling pathway molecules (e.g., *Shh, Ptch1/2, Gli1/2/3, Smo*), loss of *Tfap2* resulted in an upregulation of members in the *Fox* transcription factor family in the FNP. FOX factors normally act as downstream targets for this signaling input and functions during BA1 CNCC patterning and cartilage differentiation (Jeong et al., 2004; Xu et al., 2022; Xu et al., 2018; Xu et al., 2021). Further, hyperactivation of Hedgehog activity is strongly linked with failure of the FNP to fuse at the midline (Brugmann et al., 2010; Wu et al., 2022). Thus, it is possible that the TFAP2 midface pathway—and potentially ALX—plays a role in safeguarding FNP fusion in part by restricting *Fox* gene expression.

TFAP2 may also interact in parallel with other midfacial transcription factors. One such example includes SIX2, whose gene expression coincides with *Tfap2* in the FNP (Fig. S12) (Minoux et al., 2017) and is largely unaffected by loss of *Tfap2* in NCCs. Loss of human *SIX2* causes FND pathology (Hufnagel et al., 2016), while mice harboring a similar locus deletion produced a nasal cleft (Fogelgren et al., 2008). Mouse genetic analyses suggested SIX paralogs contribute to *Alx* gene expression (Liu et al., 2019), thereby hinting at a transcriptional battery running in parallel with TFAP2 in the FNP. Such interplay at midfacial GRN nodes requires further investigation, as it is unclear whether the SIX2 midface pathway is dependent or independent of TFAP2 activity. Given the collective contributions of both signaling pathways and transcription factors to the midface GRN, addressing these remaining knowledge gaps will further advance our understanding of how TFAP2 contributes to facial shape variation, vertebrate facial evolution, and the pathogenesis of midfacial disorders.

## MATERIALS & METHODS

### Animal husbandry and procedures

#### Mice

Experiments were all conducted in accordance with all applicable guidelines and regulations, following the “Guide for the Care and Use of Laboratory Animals of the National Institutes of Health”. The animal protocols utilized were approved by the Institutional Animal Care and Use Committee of the University of Iowa (protocols #9012197 and #2032197) and University of Colorado (protocol #14). Noon on the day of an observable copulatory plug was denoted as embryonic day 0.5 (E0.5). Additionally, all dissections were performed in 1X PBS (Fisher Scientific) treated with Diethyl Pyrocarbonate (Fisher Scientific) (i.e., DEPC-PBS). Yolk sacs or tails clips were used to extract DNA for all PCR-genotyping as previously described, with the listed primers (Table S3, Tab 1) (Van Otterloo et al., 2018). During single-cell RNA sequencing (scRNA-seq), we deployed a rapid lysis protocol using reagents from the SYBR Green Extract-N-Amp Tissue PCR Kit (Sigma) (Doan and Monuki, 2008). Yolk sacs were incubated in a 4:1 mixture of Extraction and Tissue Preparation Solutions first at room temperature (RT) for 10 min and subsequently at 95°C for 3 min. Neutralization Solution B was then added to the lysate, which was briefly vortexed prior to immediate PCR-genotyping.

Embryonic lethality was denoted for a subset of E18.5 embryos in the *Wnt1:CRE* model (see ‘Mouse alleles and breeding schemes’, Table S4), a consequence of reduced norepinephrine and compromised survival of neural crest cell (NCC)-derived tissues in the sympathetic nervous system (Schmidt et al., 2011). To prevent this lethality, pregnant dams received a water supplement to drink *ad libitum* (Hendershot et al., 2008; Lim et al., 2000). USP grade L-phenylephrine (β-adrenergic receptor agonist), isoproterenol (α-adrenergic receptor agonist), and L-ascorbic acid (agonist preservative) were added to their drinking water from E8.5 to E18.5, which was replaced every 3-5 days. Like previous studies (Abe et al., 2021), we note the supplementation did not alter craniofacial phenotypes in control or mutant embryos compared to those without supplementation.

#### Zebrafish

All work with zebrafish was approved by the University of Colorado Institutional Animal Care and Use Committee (protocol #00188). All zebrafish experiments were performed on the AB strain. Animals were maintained, staged, and genotyped as previously described (Knight et al., 2003; Mitchell et al., 2021).

### Mouse alleles and breeding schemes

*Wnt1:CRE* (Danielian et al., 1998), *Sox10:CRE* (Matsuoka et al., 2005), *Gt(ROSA)26Sortm1Sor/J* [*r26r-lacZ;* (Soriano, 1999)], and *Gt(ROSA)26Sortm4(ACTB-tdTomato,-EGFP)Luo/J* [*mTmG;* (Muzumdar et al., 2007)] mice were obtained from the Jackson Laboratory (Bar Harbor, ME). The *Tfap2a* and *Tfap2b* alleles used in this study include *Tfap2a*-null (Zhang et al., 1996), *Tfap2atm2Will/J* [*Tfap2a* floxed conditional; (Brewer et al., 2004)], *Tfap2b*-null (Martino et al., 2016), and *Tfap2btm2Will* [*Tfap2b* floxed conditional; (Van Otterloo et al., 2018)].

Females in this study were double homozygous for *Tfap2a* and *Tfap2b* floxed conditional alleles (Fig. 1A-C). For lineage tracing and fluorescence-activated cell sorting, the dams were also homozygous for *r26r-lacZ* and *mTmG* alleles, respectively. Conversely, the sires were *CRE* hemizygous and heterozygous for either *Tfap2a/b*-null alleles (used in the null/conditional model, Fig. 1A) or *Tfap2a/b* floxed conditional alleles (used in the conditional/conditional model, Fig. 1B) (Van Otterloo et al., 2018; Woodruff et al., 2021). Using either NCC-specific *Wnt1:CRE* or *Sox10:CRE* transgene, we were able to acquire *CRE*-negative controls as well as littermates that harbored NCC-specific heterozygosity for *Tfap2a* and *Tfap2b* (*Tfap2*^HET^), NCC-specific loss of *Tfap2a* (*Tfap2a*^NCKO^), NCC-specific loss of *Tfap2b* (*Tfap2b*^NCKO^), and NCC-specific loss of both *Tfap2a* and *Tfap2b* (*Tfap2*^NCKO^) (Fig. 1C). To distinguish between breeding schemes, genotype nomenclature for the null/conditional model was denoted with the “Δ/NCKO” superscript (i.e., *Tfap2a*^Δ/NCKO^, *Tfap2b*^Δ/NCKO^, *Tfap2*^Δ/NCKO^) (Fig. 1C).

### Zebrafish alleles

The transgenic line *Tg(fli1a:EGFP)y1* (Lawson and Weinstein, 2002) and the germline mutant lines for *tfap2a low*^ts213^ and *alx3*^co3003^ have been previously reported (Knight et al., 2003; Mitchell et al., 2021).

### Scanning electron microscopy

Preparation of E11.5 *CRE*-negative control, *Tfap2*^HET^, and *Tfap2*^NCKO^ littermate heads and imaging was performed as previously described (Van Otterloo et al., 2022). Briefly, samples were fixed in 4% paraformaldehyde, and then rinsed in 1X PBS. Subsequently, samples were rinsed in ddH2O, air-dried for at least 3 days, mounted on an SEM stub, and sputter coated for 4.5 minutes at 5 mA using a gold/palladium target in an Emitech K550X sputter coater. Scanning electron micrographs were acquired using a Hitachi (Tokyo, Japan) S4800 electron microscope operated in high-vacuum mode at 1.8kV. Imaging was performed through the University of Iowa Central Microscopy Research Facility workflow.

### β-galactosidase staining

E10.5 littermates with *Wnt1:CRE*-mediated recombination of the *r26r-lacZ* reporter allele were subject to β-galactosidase staining as previously described (Van Otterloo et al., 2016).

### Tissue section analyses

#### Hematoxylin and Eosin staining

Histology staining analyses of E17.5 control and *Tfap2*^Δ/NCKO^ embryos was performed as previously described (Van Otterloo et al., 2018).

#### Cryosectioning

Cryosectioning was performed as previously described (Van Otterloo et al., 2016), with minor modifications. In brief, E11.5 wild-type CD-1 mice were fixed overnight in 4% PFA at 4°C. Fixed embryos were then transferred through increasing concentrations of Tissue Tek OCT compound (Sakura Finetek USA, Inc.), embedded in 100% OCT, and stored at -80°C when not immediately processed. Embryonic heads were sectioned in a horizontal fashion (10 µm slices) using a Microm HM525 NX Cryostat Microtome (ThermoFisher) housed within the University of Iowa Central Microscopy Research Facility, mounted onto positively charged glass slides, and were left in a closed container at RT overnight to further adhere. We used the eMouseAtlas database (Armit et al., 2017) to reference the horizontal section angle.

#### Immunofluorescence

Sectioned tissue was blocked for 1.5 hours in 5% milk or 3% BSA dissolved in 0.1% Triton-X/PBS (PBSTx). Wild-type sections were used to examine protein expression of TFAP2A (3B5-supernatant, Santa Cruz Biotechnology; 1:25 dilution) and TFAP2B (2509S, Cell Signaling Technology; 1:25 dilution). After washing with PBSTx, samples were incubated with Alexa Fluor 488/568 secondary antibodies (Invitrogen; 1:500 dilution) for 1.5 hours. We used DAPI to counterstain nuclei. Stains were imaged via Zen software on a Zeiss 700 LSM confocal microscope. Images were exported as Z-stacked, maximum intensity projection TIFF images.

### Micro-computed tomography

The University of Iowa Small Animal Imaging Core Facility provided scanning and software services. E12.5 *CRE*-negative control and *Tfap2*^NCKO^ littermates were prepared as previously described (Hsu et al., 2016). All fixing, washing, and incubation steps were performed at 4°C. In brief, embryos were fixed in 4% PFA overnight and washed thrice and stored in 1X PBS until the day of scanning was determined. Three days prior to scanning, embryos were incubated in 1 N (v/v) iodine solution (Sigma). Control and mutant embryos were then mounted above 1% agarose in the same screw-top conical tube, submerged in 1X PBS, and immediately imaged at 5- or 6-µm resolution.

Sectional analysis and measurements of the µCT scans were performed on Dragonfly software (Object Research Systems). Embryo scans were oriented to a frontal plane to examine nasal morphologies. Embryos were also oriented in a horizontal fashion, spanning from the nasal pits to the hindbrain to quantify the distance between the anterior tips of the mFNP. Control and mutant pairs were imaged using the same parameters. A Student’s t-test was performed on GraphPad Prism using three independent control-mutant pairs to compare mFNP tip distances. A p-value less than 0.05 was considered statistically significant.

### Concurrent Alizarin Red and Alcian Blue staining preparation

Concurrent Alizarin Red/Alcian Blue staining of E18.5 embryos and E15.5 Alcian Blue staining of null/conditional embryos were performed as previously described (Van Otterloo et al., 2016). Note that Alcian blue staining for E18.5 nasal capsule elements are highly variable.

### Bulk RNA-sequencing

#### Tissue preparation, cDNA library generation, and sequencing

E10.5 FNP and MxP bulk tissue from three control and three *Tfap2*^Δ/NCKO^ littermates were processed for RNA-seq as previously described (Van Otterloo et al., 2016). After PCR-genotyping, samples stored in RNAlater (ThermoFisher) were processed for RNA (Norgen Biotek) and further purified with the RNAeasy kit (Qiagen). mRNA quality was determined with DNA Analysis ScreenTape (Agilent Technologies), and cDNA libraries were generated using the Illumina TruSeq Stranded mRNA Sample Prep Kit (Illumina). Samples were sequenced on the Illumina HiSeq2500 platform as 150-bp single-end reads. Library construction and sequencing was carried out by the Genomics and Microarray Core on the University of Colorado Anschutz Medical Campus.

#### RNA-seq dataset processing and analyses

Following sequencing, reads were demultiplexed, and FASTQ files processed using two distinct bioinformatic pipelines. Two pipelines were used to help reduce false positives. In the first approach, FASTQ files were mapped to the mm10 genome using STAR Aligner (Dobin et al., 2013) with default settings. Gene expression was then quantified using StringTie (Pertea et al., 2016; Pertea et al., 2015), with the following settings: -e (expression estimation mode), -G (mm10 reference annotation file), -A (gene abundances). Following quantification, count matrices were produced using the *prepDE.py* script, available from StringTie, and used as input for DESeq2 (Love et al., 2014). Differential gene expression analysis was conducted in DESeq2 using the *DESeqDataSetFromMatrix* function. For the second approach, FASTQ files were pseudo-aligned to the mm10 genome using the kallisto (Bray et al., 2016) *quant* function with the following settings: -b 50, --single, -l 100, -s 20. After alignment, differential gene expression analysis was conducted using sleuth (Pimentel et al., 2017), along with the *sleuth_prep* function and the following settings: full_model = genotype, gene_model = TRUE, read_bootstrap_tpm = TRUE, extra_bootstrap_summary = TRUE, transformation_function = function(x) log2(x + 0.05)). The resulting ‘sleuth object’ was then subjected to the *sleuth_fit* function, followed by the *sleuth_wt* function to calculate differentially expressed genes between genotypes. The kallisto-sleuth pipeline output was used, along with ggplot2 (Wickham, 2016) and ggrepel, to generate and visualize differentially expressed genes by volcano plot. Genes that both had, 1) a fold-change greater or less than +1.25 or -1.25, respectively, and an adjusted p-value less than 0.05 (pipeline 1), and 2) a q-value less than 0.1 (pipeline 2) (Table S1), were used for ontology, enrichment, and pathway analysis using the Enrichr platform (Chen et al., 2013; Kuleshov et al., 2016; Xie et al., 2021).

#### Reanalysis and visualization of published RNA-seq datasets

To directly compare the E11.5 mandibular and frontonasal prominence mesenchyme (e.g., Fig. 2) a procedure like ‘pipeline 2’ above (kallisto and sleuth) was used. Specifically, triplicate bam files were downloaded from the Facebase.org repository (FB00000867) for both E11.5 mandibular prominence mesenchyme (Biosample 30H6; RID 33SA, 33SE, 33SJ) and E11.5 frontonasal prominence mesenchyme (Biosample 2Y6P; RID: 33TT, 33TY, 33V2). Next, bam files were converted to fastq files using the *bam2fq* function in samtools (Li et al., 2009), Generated FASTQ files were then used along with kallisto and sleuth for differential gene expression analysis. Data was plotted using ggplot2 and ggrepel.

### Mouse single-cell RNA-sequencing

#### Single-cell dissociation and cell sorting

To isolate CNCCs, E11.5 embryonic heads were cut just below BA2 and the otic vesicle by microdissection. Using *Wnt1:CRE*-mediated recombination of the *mTmG* reporter allele (Muzumdar et al., 2007), *CRE*-positive embryos were selected for further processing. Tissue samples were then subject to a cold protease single-cell dissociation protocol (Adam et al., 2017), with modifications. Embryo heads were incubated in a protease cocktail comprised of *Bacillus licheniformis* Subtilisin A (Creative Enzymes), Accumax, and Accutase (Fisher Scientific) suspended in DEPC-PBS as previously described (Sekiguchi and Hauser, 2019). The tissue was then carefully disrupted using a disposable Eppendorf pellet pestle and placed on a rotator at 4°C for 30 min. During this step, samples were gently triturated every 10 minutes using a wide bore 1 mL pipette tip. The enzymatic reaction was quenched using an equal volume of 10% heat-inactivated FBS (Fisher Scientific) in 1X phenol red-free DMEM (ThermoFisher). Dissociated cells were washed thrice in DEPC-PBS by centrifugation at 600 × g at 4°C for 5 min and then transferred, through Bel Art SP Scienceware Flowmi 40-μm strainers (ThermoFisher), into chilled flow tubes precoated with DEPC-PBS. After GFP-positive sorting on the University of Iowa Flow Cytometry Facility’s ARIA II system (Becton Dickinson), CNCCs were re-suspended in DEPC-PBS containing 0.04% non-acetylated BSA (New England Biolabs). Cell viability was determined to be >95% by Trypan Blue staining. During cell sorting, genotypes were confirmed by rapid genotyping, allowing isolation of GFP-positive cells from two *Tfap2*^HET^ control and two sibling-matched *Tfap2*^NCKO^ mutant embryos. Cells from each “biological replicate” were combined to generate a single control and mutant sample.

#### Library preparation & sequencin

∼6,000 GFP-positive CNCCs from each condition were subject to library preparation by the University of Iowa Genomics Division on the 3’ expression scRNA-seq 10X Chromium v3.1 pipeline. Libraries were sequenced on an Illumina NovaSeq 6000 platform as 100-bp paired-end reads. Over 20,000 reads per cell were acquired, resulting in ∼1.5 × 10^8^ reads per condition.

#### Quality control, integration, and UMAP visualization

Sequencing results were demultiplexed and converted to a FASTQ format using the Illumina bcl2fastq software. Reads were processed and aligned to the mm10 reference genome assembly with the Cell Ranger *Count* function. Seurat v4.3.0 was used for quality control, filtering out cells with >10% mitochondrial counts, less than 200 features, and more than 7,500 features. We then integrated *Tfap2*^HET^ and *Tfap2*^NCKO^ conditions (Hao et al., 2021), where RNA counts were normalized and *FindVariableFeatures* was run with the following parameters: selection.method = “vst”, nfeatures = 2,000. Cell clustering by Uniform Manifold Approximation and Projection (UMAP) plots was deployed by *RunPCA* and then *RunUMAP* with dims 1:30. The UMAP visualizing all CNCCs was generated with *FindClusters* using a resolution of 0.40. Major CNCC derivatives were annotated based on published sectional immunofluorescence and scRNA-seq profiling (Soldatov et al., 2019). Gene expression was plotted on UMAP plots by *FeaturePlots*. Gene expression mapping of CNCC lineage markers was done using the *FeaturePlot* function.

#### Cell cycle analysis

The *CellCycleScoring* function was performed as previously described, with minor modifications (Hao et al., 2021). The UMAP resolution was changed to 0.05 to segregate the major CNCC lineages as individual clusters. Bioconductor BiomaRt v2.46.3 (Durinck et al., 2005) was used to convert a published list of cell cycle genes (Kowalczyk et al., 2015) from human to mouse gene nomenclature. Cell cycle scores were then mapped directly onto the UMAP and graphed, as a percentage relative to each CNCC lineage, using GraphPad Prism.

#### Cellular distribution analysis

To examine genotype cellular distributions in each cluster, each condition was expressed as a percentage per cluster. These analyses were visualized in GraphPad Prism. For the initial UMAP containing all sorted cells, a resolution of 0.05 was used. For the MAGIC-clustered UMAP, a resolution of 0.10 was used (see “Facial prominence mesenchyme annotation”).

#### Facial prominence mesenchyme annotation

To better define the facial prominence subpopulations, we subset and re-clustered the “Mesenchyme” clusters using MAGIC v2.0.3 (van Dijk et al., 2018) with *RunUMAP* dims 1:20. Major mesenchyme populations were then annotated based on known gene markers, which were plotted on the MAGIC-clustered UMAP with DefaultAssay = “MAGIC_RNA”. A cluster resolution of 0.10 was chosen to best visualize each prominence population and the second branchial arch as individual clusters based on published *in situ* data, ChIP-seq, ATAC-seq, and transcriptomic datasets (Gu et al., 2022; Hooper et al., 2020; Minoux et al., 2017). To visualize co-expression of two genes on the UMAP, the flag *blend = “TRUE”* was used in the *FeaturePlots* function.

#### Gene-set and gene expression analyses

The UMAP resolution was set to 0.05 to distinguish the major cell groupings, to which we then subset the “pseudobulked” mesenchyme group. We then used the *FindMarkers* function with ident.1 = *Tfap2*^HET^ and ident.2 = *Tfap2*^NCKO^ on CNCCs, to generate a list of differentially expressed genes (adjusted p-value < 0.05 and a log fold-change threshold of 0.10) (Table S2, Tab 9). For gene-set enrichment, ontology, and pathway analysis, lists of either up- or downregulated genes, generated from the mesenchyme “peudobulk” approach, in the *Tfap2*^NCKO^ condition were inputted in the Enrichr pipeline. Control-versus-mutant expression analysis of select genes was visualized as a violin plot using the *VlnPlot* function on the MAGIC-clustered UMAP, with DefaultAssay = “RNA”. Differential gene expression was validated for a select set of genes-of-interest by targeted approaches (see real-time PCR).

#### Gene-set overlap with bulk RNA-seq datasets

First, to determine overlapping genes between the E11.5 scRNA-seq and E10.5 bulk RNA-seq datasets, gene lists of differentially expressed genes were compiled (Table S1, Tab 11). Datasets used included the following outputs: E11.5 scRNA-seq mesenchyme “pseudobulk” (Table S1, Tab 10), E10.5 bulk FNP/MxP (i.e., ‘upper-face’) pipeline 1 (Table S1, Tab 4) and pipeline 2 (Table S1, Tab 7), and E10.5 bulk MdP (i.e., ‘lower-face’) pipeline 1 and pipeline 2 (Van Otterloo et al., 2018). Second, the VLOOKUP function in Excel was used to intersect all 4 E10.5 gene lists with the E11.5 pseudobulk gene list and generate a comprehensive overlap table (Table S1, Tab 2).

### Real-time PCR

For the null/conditional model, E10.5 bulk FNP tissue was acquired from *CRE*-negative controls and *Tfap2*^Δ/NCKO^ mutants by micro-dissection. For the conditional/conditional model, E11.5 MxP and FNP tissue from *Tfap2*^HET^ and *Tfap2*^NCKO^ embryos were microdissected, and CNCCs were collected by the *Wnt1:CRE-mTmG* sorting strategy. For the latter, single-cell dissociation was conducted using 0.25% trypsin (ThermoFisher) for 15 minutes at 37°C. Samples were either stored in RNAlater (ThermoFisher) at -20°C or immediately processed for RNA using the RNeasy Mini Kit as per manufacturer’s protocol (Qiagen). Real-time analyses were performed as previously described (Van Otterloo et al., 2018). After PCR-genotyping, cDNA was prepared with SuperScript III First-Strand Synthesis Kit (Invitrogen/ThermoFisher), where equal amounts of RNA template were processed from each condition. Real-time reactions were performed on a CFX Connect instrument (Bio-Rad) with either SYBR Green or Select Master Mixes (Applied Biosystems, ThermoFisher). Primers targeting *Alx1*, *Alx3*, and *Alx4* transcripts were designed to target exons flanking intronic sequences. Relative mRNA expression levels were normalized to *B2m* or *Actb* transcripts. Real-time PCR primers are listed in Table S3, Tab 2.

### Whole mount *in situ* hybridization

Control and *Tfap2*^Δ/NCKO^ embryos at E9.0 and E10.5 were subject to whole mount *in situ* hybridization with *Alx3* and *Alx4* RNA probes as previously described (Van Otterloo et al., 2018). Primers used to generate the probes are listed in Table S3, Tab 3.

### Chromatin Immunoprecipitation followed by sequencing

#### Tissue preparation, library construction, and sequencing

Chromatin immunoprecipitation (ChIP) was conducted, essentially as previously described (Van Otterloo et al., 2022), with the exception that full facial prominences were used, rather than just isolated ectoderm. Further modifications included the use of an anti-TFAP2A antibody (5 µg, sc-184, Santa Cruz Biotechnology) and Dynabeads (ThermoFisher Scientific). Following precipitation and subsequent quality control analysis (ScreenTape analysis, Agilent), samples (i.e., sc-184 IP and input DNA) were sequenced on the Illumina HiSeq 2500 platform as 150-bp single-end reads. Library construction and sequencing was carried out by the Genomics and Microarray Core at the University of Colorado Anschutz Medical Campus.

#### Peak calling and analyses

Following sequencing and demultiplexing, FASTQ files were mapped to the mm10 genome using BWA (Li and Durbin, 2009), followed by sam to bam file conversion using Samtools (Li et al., 2009). To identify ‘enriched’ regions (i.e., regions with significantly more reads in the TFAP2 IP versus input), MACS2 was utilized with the following settings: -f BAM, -g mm, --keep-dup 2 (Table S2, Tab 3). Motif and peak annotation were completed using the *findMotifsGenome.pl* and the *annotatePeaks.pl* functions in HOMER (Heinz et al., 2010). Heatmap visualization of enriched regions was completed using the *bamCoverage*, *computeMatrix* and *plotHeatmap* function in deepTools (Ramirez et al., 2016), with the following settings: -of ‘bigwig’, --ignoreDuplicates, -bs 25, --normalizeUsing RPKM (for *bamCoverage*); reference-point, -b 3000, -a 3000 –referencePoint center (for *computeMatrix*); -- whatToShow ‘heatmap and colorbar’, --missingDataColor 1, --kmeans 7, --refPointLabel “peak” (for *plotHeatmap*). For *computeMatrix* (Fig. 5E) the following scaled bigwig datasets were downloaded from GEO (Barrett et al., 2013) and used (along with the MACS2 generated ‘TFAP2-bound’ bed file): GSM2371708, E10.5 FNP H3K4me2; GSM2371717, E10.5 FNP H3K27ac; GSM2371732, E10.5 FNP ATAC-seq; GSM2371713, E10.5 FNP H3K27me3 (peak-cluster annotations in Table S2, Tab 3) (Minoux et al., 2017). Additional datasets used for analysis included: GSM2371710, E10.5 MdP H3K4me2; GSM2371719, E10.5 MdP H3K27ac; GSM2371734, E10.5 MdP ATAC-seq; GSM2371715, E10.5 MdP H3K27me3. The Integrative Genomics Viewer (Robinson et al., 2011) was used for gene level visualization of these datasets at *Alx* loci. Pathway analysis was conducted using GREAT (McLean et al., 2010) with the following setting: ‘*Basal plus extension*’, ‘Proximal 5 kb upstream, 1 kb downstream, plus Distal: up to 100 kb’. To intersect peak coordinates with dysregulated genes, a data matrix was generated (Table S2, Tab 11) that included all genes in the GREAT database (mm10, n = 20,510), their TFAP2 peak association (yes/no) using the criteria outlined above, along with gene lists from the E10.5 bulk RNA-seq analysis and the E11.5 mesenchyme “pseudobulk” analysis. The E10.5 bulk RNA-seq datasets included the ‘upper-face’ pipeline 1, pipeline 2, and their overlap (this study) along with the ‘lower-face’ pipeline 1, pipeline 2, and their overlap (Van Otterloo et al., 2018). Total genes with or without peaks (excluding those that did not have a 1:1 match between datasets) were summed and summarized (Table S2, Tab 2). To correlate peak status with gene expression in the midface, genes with and without an associated TFAP2 peak were paired with previously calculated FPKM values from the E11.5 FNP mesenchyme (Hooper et al., 2020) (Table S2, Tab 12). Data was plotted and visualized using ggplot2 and statistical significance calculated using ggpubr.

### Zebrafish single-cell RNA sequencing analysis

Expression mapping on UMAPs of published zebrafish CNCC scRNA-seq datasets were performed as previously described (Stenzel et al., 2022).

### *in situ* hybridization chain reaction v3.0 and fluorescence microscopy

We ordered oligos from Molecular Instruments targeting the same *alx3* 5’ UTR sequence previously used for *in situ* hybridization (Mitchell et al., 2021) and followed the manufacturer’s protocol for whole-mount zebrafish embryos and larvae (Choi et al., 2016). High salt content in buffers used during the hybridization chain reaction (HCR) v3.0 protocol made genotyping difficult. To circumvent this, we used Promega GoTaq Flexi (M8295) in “M” buffer (2mM MgCl2, 14mM Tris-HCl pH 8.4, 68.25 mM KCl, 0.0013% gelatin, 1.8 mg/mL BSA, 140 µM each dNTP). Embryos or larvae were then embedded in 0.2% agarose and imaged with an Andor Dragonfly 301 spinning disk confocal system. Acquisition parameters and fluorescence adjustments were applied linearly and equally to all samples.

### Zebrafish genetic interaction analyses

Zebrafish larvae 6 days post-fertilization were subject to concurrent Alizarin Red/Alcian Blue staining preparations as previously described (Walker and Kimmel, 2007). Nomarski imaging, genotype-blinded phenotype scoring, and statistical analysis were performed as previously described (Bailon-Zambrano et al., 2022; Mitchell et al., 2021; Sucharov et al., 2019). In brief, larval skeletons were dissected and flat mounted for imaging on a Leica DMi8 inverted microscope equipped with a Leica DMC2900. For penetrance scores, we performed the Fisher’s exact test using GraphPad. All sample sizes and p-values are included in Fig. 6F.

## FOOTNOTES

## Acknowledgements

The authors would like to acknowledge the University of Iowa Small Animal Imaging Core, Central Microscopy Research Facility, Flow Cytometry Facility, and Genomics Division for their technical support in µCT, tissue processing, and scRNA-seq experiments. These facilities are funded through user fees and generous financial support from the Carver College of Medicine. To Dr. Michael Chimenti from the Bioinformatics Division, Professor Huojun Cao from the College of Dentistry Division of Biostatistics and Computational Biology, Nate Mullin, and Yann Vanrobaeys for their insights into scRNA-seq analyses. To Jamie Thompson for technical input on immunofluorescence experiments and additional care for animals used in this study. Lastly, we are grateful to all Van Otterloo lab members and our colleagues in the Craniofacial Interest Group for their invaluable feedback, particularly Professors Colin Kenny, Robert Cornell, Brad Amendt, Martine Dunnwald, and John Manak.

## Author Contributions

<u>Conceptualization</u>: TTN, JMM, TJW, JTN, EVO

<u>Methodology</u>: TTN, JMM, TJW, JTN, EVO

<u>Software</u>: TTN, JMM, KLJ, EVO

<u>Validation</u>: TTN, JMM, EVO

<u>Formal analysis</u>: TTN, JMM, JTN, EVO

<u>Investigation</u>: TTN, JMM, MDK, EVO

<u>Resources</u>: KLJ, TW, JTN, EVO

<u>Data Curation</u>: TTN, JMM, EVO

<u>Writing – original draft</u>: TTN, JMM

<u>Writing – review and editing</u>: TTN, JMM, TJW, JTN, EVO

<u>Visualization</u>: TTN, JMM, EVO

<u>Supervision</u>: TJW, JTN, EVO

<u>Project administration</u>: TJW, JTN, EVO

<u>Funding acquisition</u>: TTN, TJW, JTN, EVO

## Summary Statement

Mouse and zebrafish genetic analyses and next-generation sequencing profiling identify a critical, conserved TFAP2-*ALX* transcriptional axis within the midface developmental gene regulatory network.

## Competing Interest Statement

The authors declare no conflict of interest.

**Fig. S1 – Histological analysis of midface phenotypes in *Tfap2*^Δ/NCKO^ mutants.** (**A-F**) Coronal sections of H&E stained anterior (A-C) and posterior (D-F) regions of the snout of E17.5 embryos, indicated by genotype. Note, the schematic in the top left highlights relative position of sections in A-C (‘Top’) and D-F (‘Bottom’). Yellow lines in panels C and F demarcate the midfacial cleft. Scale bars = 500 µm. N = 3 per genotype. Abbreviations: i, incisor; ns, nasal septum; t, tongue; v, vibrissae.

**Fig. S2 – TFAP2A and TFAP2B cooperative function in CNCCs during midface fusion.** (**A-C**) Gross craniofacial morphology, as detected by scanning electron microscopy (A) or brightfield imaging (B, C) of E11.5 (A), E12.5 (B), or E15.5 (C) embryonic midfaces, as indicated by genotype. Panel A is ventral views, whereas panels B and C are top-down views of the cranium and midface. Black dotted lines in the E12.5 images highlight the frontonasal prominences. White arrowheads point to the midface cleft. N = 3 per genotype. Scale bar = 1 mm. Abbreviations: FNP, frontonasal prominence; lFNP, lateral FNP; mFNP, medial FNP; np, nasal pit.

**Fig. S3 – *Tfap2*^NCKO^ mutant skeletal phenotypes recapitulate those observed in *Tfap2*^Δ/NCKO^ mutants.** (**A**) An E18.5 *Wnt1:CRE Tfap2*^NCKO^ skeletal preparation showing ‘maxilla-like’ medial projections on the mandible that meet at the midline (left) along with syngnathia of the left jaw viewed laterally in isolation (right). (**B**) Isolated left mandible from E18.5 *Tfap2*^NCKO^ embryos, with indicated genotype. These phenotypes match those previously described in *Tfap2*^Δ/NCKO^ embryos (Van Otterloo et al., 2018). Scale bar = 1 mm. Abbreviations: agp, angular process; cdp, condylar process; crp, coronoid process; fpm, frontal process of the maxilla; jg, jugal bone; li, lower incisor; mc, Meckel’s cartilage; mx-l, maxilla-like; ppmx, palatal process of the maxilla; zpm, zygomatic process of the maxilla.

**Fig. S4 – Lineage tracing and immunofluorescence analysis suggest a post-migratory role for TFAP2A and TFAP2B during midfacial development.** (**A**) Front view of E10.5 embryos, with indicated genotypes. Cranial neural crest cells are β-galactosidase-stained from *Wnt1:CRE*-mediated recombination of the *r26r-lacZ* reporter allele. Dashed yellow lines mark the midline-proximal edges of the medial domains of the frontonasal prominences (mFNP). White arrowhead in Tfap2^NCKO^ embryo points to the increased gap between the mFNP. Scale bar = 500 µm. (**B**) Co-staining of nuclei (DAPI, blue), TFAP2A (gold), and TFAP2B (red) in midface tissue. The top-left image, derived from BioRender, indicates the plane of each section that TFAP2 co-staining was performed. The bottom-left image, derived from the eMouse atlas (Armit et al., 2017), is a representative orientation of the tissue section. Boxed is the region examined in the immunofluorescent images. The white arrowhead indicates absent TFAP2B protein expression in the nasal epithelium, in contrast to the top panel. White dashed lines indicate boundaries between mesenchyme and epithelium. Additional abbreviations: d, dorsal; ect, ectoderm; lFNP, lateral domain of the FNP; MxP, maxillary prominence; ne, nasal epithelium; np, nasal pit; v, ventral.

**Fig. S5 – A post-migratory CNCC role for TFAP2A and TFAP2B during midfacial development.** Frontal view of E18.5 *Sox10:CRE*-*Tfap2^NCKO^* animals, with indicated genotypes. Scale bar = 1 mm.

**Fig. S6 – Defects in the midfacial skeleton of *Sox10:CRE* Tfap2^NCKO^ mutants.** (**A-A”, B-B”**) Three individual E18.5 *Sox10:CRE*-*Tfap2*^NCKO^ skeletal preparations in top-down (A) or bottom-up (B) view. Anterior is to the left. Yellow dashed lines in the top-down view outline the peripheral edges of the calvarial bones while those in the bottom-up view outline the presphenoid bone. Yellow arrowheads point to the cartilaginous ectopias stemming from the nasal septum. Note, in contrast to the first two embryos, the maxillary bone has been removed from the third. Further, the third exhibits isolated Alizarin red stained tissue (i.e., bony ‘islands’, individually outlined) on the calvaria instead of the cartilaginous ectopias. Scale bar = 1 mm. Abbreviations: bs, basisphenoid; cs, coronal suture; fr, frontal bone; ip, interpareital bone; nl, nasal/ethmoid labyrinth; ns, nasal septum; pl, palatine; pmx, premaxilla; ppp, palatal process of the palatine; ppmx, palatal process fo the premaxilla; ppro, pila postoptica; ppso, pila postoptica; pr, pareital bone; vm, vomer bone.

**Fig. S7 – Jaw phenotypes in Sox10:CRE-Tfap2NCKO mutants partially recapitulate those observed in Wnt1:CRE-Tfap2NCKO and -Tfap2Δ/NCKO mutants.** (**A-F**) E18.5 maxillary and mandibular bones in isolation, with indicated genotypes. In panels A through D, the left-side elements are presented; panels E and F show the right-side elements. In panel B, circled is the missing jugal bone and thicked zygomatic process of the maxilla. In panel E, the sygnanthia is circled. In panel F, fusion between the the jugal and thickened zygomatic process of the maxilla is circled. Scale bar = 1 mm. Abbreviations: agp, angular process; cdp, condylar process; crp, coronoid process; fpm, frontal process of the maxilla; jg, jugal bone; li, lower incisor; mc, Meckel’s cartilage; ppmx, palatal process of the maxilla; sq, squamosal bone; zpm, zygomatic process of the maxilla.

**Fig. S8 – Wnt1:CRE-Tfap2^Δ/NCKO^ mutants present defects in the midfacial skeleton.** (**A**) Oblique frontal view of Alcian Blue stained preparations for E15.5 *Tfap2*^Δ/NCKO^ mutant embryos, indicated by genotypes. The midface is outlined by black dotted lines. The black arrowhead points to ectopic cartilage forming superior to the midface cleft. (**A’**) Bottom-up ventral view, anterior to the left, of chondrocrania, as in panel A, with a focus on nasal cartilage elements. Black arrowheads point to structural aberrations which presumably would have formed the anterior nasal capsules and labyrinths. (**B**) Top-down view of E18.5 craniums, anterior up, from the indicated genotypes. Gold dashed lines outline the edge of the calvarial bones at the midline. The asterisk marks the midface cleft and missing nasal bones. N = 3 per genotype. Scale bar = 500 µm. Abbreviations: cs, coronal suture; e, eye; fr, frontal bone; ip, interparietal bone; ms, metopic suture; na, nasal bone; ns, nasal septum; pn, paries nasi; ppro, pila preoptica; ppso, pila postoptica; pr, parietal bone; ss, sagittal suture.

**Fig. S9 – Gene expression-based scRNA-seq annotation of the major CNCC lineages.** Gene expression of select genes mapped onto the Uniform Manifold Approximation and Projection (UMAP) plot in Fig. 4C. These genes were selected based on published scRNA-seq datasets (Soldatov et al., 2019). Note that cells from the brain are included in this dataset because *Wnt1:CRE* labels parts of the brain.

**Fig. S10 – Cellular profiling of the three major CNCC lineages in the developing head.** (**A**) *Tfap2*^HET^ and *Tfap2*^NCKO^ representation in the integrated dataset, divided by the three major groupings and expressed as a percentage. The dashed line marks the fifty-percent threshold. (**B**) Gene expression-based cell cycle scoring for indiivdual *Tfap2*^HET^ and *Tfap2*^NCKO^ conditions. Cell cycle scores are mapped on the Uniform Manifold Approximation and Projection (UMAP) plots of individual conditions (left) and visualized as percentages per CNCC lineage.

**Fig. S11 – Enrichment analyses of downregulated genes in the scRNA-seq mesenchyme group.** Terms from the Enrichr pipeline (Kuleshov et al., 2016), based on (**A**) downregulated or (**B**) upregulated genes identified through the mesenchyme psuedobulk analysis (Table S2, Tab 9).

**Fig. S12 – Gene expression-based scRNA-seq annotation of the major CNCC mesenchymal populations.** MAGIC-based (van Dijk et al., 2018) gene expression of select genes mapped onto the Uniform Manifold Approximation and Projection (UMAP) plot in Fig. 4D. These genes were selected based on published transcriptomic datasets (Gu et al., 2022; Hooper et al., 2020), epigenome datasets (Minoux et al., 2017), and published *in situ* data.

**Fig. S13 – *Tfap2* and *Alx* paralog transcripts are enriched in the murine FNP CNCCs.** Gene expression of *Tfap2a* and *Tfap2b* (blue) overlaid with individual *Alx* paralogs (red) onto the Uniform Manifold Approximation and Projection (UMAP) plot. Color saturation correlates to gene expression levels, while degree of co-expression is read out as a blending of the two colors.

**Fig. S14 – Cellular and gene expression profiling in the CNCC-derived mesenchyme.** (**A**) Cellular distributions of *Tfap2*^HET^ and *Tfap2*^NCKO^ conditions as Uniform Manifold Approximation and Projection (UMAP) plots (A) or as a percentage in each cluster (A’). (**B**) Violin expression plots of branchial arch (BA)-enriched genes previously identified (Van Otterloo et al., 2018). Boxed are clusters for the maxillary prominence (MxP), mandibular prominence (MdP), and BA2. (**C**) Violin expression plots for genes dysregulated in the midface. Boxed are the frontonasal prominence (FNP) clusters.

**Fig. S15 – Additional GREAT pathway analysis on TFAP2 genomic binding.** Terms enriched in cluster 3 and 4 peaks that contained positive TFAP2 ChIP-seq signal. Note, peaks in these clusters are significantly enriched for skin (cluster 4), neuronal (clusters 3, 4), and cardiac (cluster 3) terms.

**Fig. S16 – *tfap2a* and *alx3* transcripts are enriched in the zebrafish frontonasal CNCCs.** Gene expression of *tfap2a* and *alx3* overlaid together on a Uniform Manifold Approximation and Projection (UMAP) plot of a published single-cell RNA-seq dataset generated from sorted zebrafish cranial neural crest cells (Stenzel et al., 2022). Color saturation correlates to gene expression levels, while degree of co-expression is read out as a blending of the two colors.

## REFERENCES

Abe, M., Cox, T. C., Firulli, A. B., Kanai, S. M., Dahlka, J., Lim, K. C., Engel, J. D. and Clouthier, D. E. (2021). GATA3 is essential for separating patterning domains during facial morphogenesis. Development 148.

Adam, M., Potter, A. S. and Potter, S. S. (2017). Psychrophilic proteases dramatically reduce single-cell RNA-seq artifacts: a molecular atlas of kidney development. Development 144, 3625–3632.

Armit, C., Richardson, L., Venkataraman, S., Graham, L., Burton, N., Hill, B., Yang, Y. and Baldock, R. A. (2017). eMouseAtlas: An atlas-based resource for understanding mammalian embryogenesis. Dev Biol 423, 1–11.

Bailon-Zambrano, R., Sucharov, J., Mumme-Monheit, A., Murry, M., Stenzel, A., Pulvino, A. T., Mitchell, J. M., Colborn, K. L. and Nichols, J. T. (2022). Variable paralog expression underlies phenotype variation. Elife 11.

Barrallo-Gimeno, A., Holzschuh, J., Driever, W. and Knapik, E. W. (2004). Neural crest survival and differentiation in zebrafish depends on mont blanc/tfap2a gene function. Development 131, 1463–1477.

Barrett, T., Wilhite, S. E., Ledoux, P., Evangelista, C., Kim, I. F., Tomashevsky, M., Marshall, K. A., Phillippy, K. H., Sherman, P. M., Holko, M., et al. (2013). NCBI GEO: archive for functional genomics data sets--update. Nucleic Acids Res 41, D991–995.

Beverdam, A., Brouwer, A., Reijnen, M., Korving, J. and Meijlink, F. (2001). Severe nasal clefting and abnormal embryonic apoptosis in Alx3/Alx4 double mutant mice. Development 128, 3975–3986.

Bhatt, S., Diaz, R. and Trainor, P. A. (2013). Signals and switches in Mammalian neural crest cell differentiation. Cold Spring Harb Perspect Biol 5.

Brandon, A. A., Almeida, D. and Powder, K. E. (2022). Neural crest cells as a source of microevolutionary variation. Semin Cell Dev Biol.

Bray, N. L., Pimentel, H., Melsted, P. and Pachter, L. (2016). Near-optimal probabilistic RNA-seq quantification. Nat Biotechnol 34, 525–527.

Brewer, S., Feng, W., Huang, J., Sullivan, S. and Williams, T. (2004). Wnt1-Cre-mediated deletion of AP-2alpha causes multiple neural crest-related defects. Dev Biol 267, 135–152.

Brugmann, S. A., Allen, N. C., James, A. W., Mekonnen, Z., Madan, E. and Helms, J. A. (2010). A primary cilia-dependent etiology for midline facial disorders. Hum Mol Genet 19, 1577–1592.

Chen, E. Y., Tan, C. M., Kou, Y., Duan, Q., Wang, Z., Meirelles, G. V., Clark, N. R. and Ma’ayan, A. (2013). Enrichr: interactive and collaborative HTML5 gene list enrichment analysis tool. BMC Bioinformatics 14, 128.

Choi, H. M., Calvert, C. R., Husain, N., Huss, D., Barsi, J. C., Deverman, B. E., Hunter, R. C., Kato, M., Lee, S. M., Abelin, A. C., et al. (2016). Mapping a multiplexed zoo of mRNA expression. Development 143, 3632–3637.

Claes, P., Roosenboom, J., White, J. D., Swigut, T., Sero, D., Li, J., Lee, M. K., Zaidi, A., Mattern, B. C., Liebowitz, C., et al. (2018). Genome-wide mapping of global-to-local genetic effects on human facial shape. Nat Genet 50, 414–423.

Clouthier, D. E., Garcia, E. and Schilling, T. F. (2010). Regulation of facial morphogenesis by endothelin signaling: insights from mice and fish. Am J Med Genet A 152A, 2962–2973.

Cox, S. G., Kim, H., Garnett, A. T., Medeiros, D. M., An, W. and Crump, J. G. (2012). An essential role of variant histone H3.3 for ectomesenchyme potential of the cranial neural crest. PLoS Genet 8, e1002938.

Danielian, P. S., Muccino, D., Rowitch, D. H., Michael, S. K. and McMahon, A. P. (1998). Modification of gene activity in mouse embryos in utero by a tamoxifen-inducible form of Cre recombinase. Curr Biol 8, 1323–1326.

Dash, S. and Trainor, P. A. (2020). The development, patterning and evolution of neural crest cell differentiation into cartilage and bone. Bone 137, 115409.

de Croze, N., Maczkowiak, F. and Monsoro-Burq, A. H. (2011). Reiterative AP2a activity controls sequential steps in the neural crest gene regulatory network. Proc Natl Acad Sci U S A 108, 155–160.

Debbache, J., Parfejevs, V. and Sommer, L. (2018). Cre-driver lines used for genetic fate mapping of neural crest cells in the mouse: An overview. Genesis 56, e23105.

Doan, L. and Monuki, E. S. (2008). Rapid genotyping of mouse tissue using Sigma’s Extract-N-Amp Tissue PCR Kit. J Vis Exp.

Dobin, A., Davis, C. A., Schlesinger, F., Drenkow, J., Zaleski, C., Jha, S., Batut, P., Chaisson, M. and Gingeras, T. R. (2013). STAR: ultrafast universal RNA-seq aligner. Bioinformatics 29, 15–21.

Dooley, C. M., Wali, N., Sealy, I. M., White, R. J., Stemple, D. L., Collins, J. E. and Busch-Nentwich, E. M. (2019). The gene regulatory basis of genetic compensation during neural crest induction. PLoS Genet 15, e1008213

Durinck, S., Moreau, Y., Kasprzyk, A., Davis, S., De Moor, B., Brazma, A. and Huber, W. (2005). BioMart and Bioconductor: a powerful link between biological databases and microarray data analysis. Bioinformatics 21, 3439–3440.

Eckert, D., Buhl, S., Weber, S., Jager, R. and Schorle, H. (2005). The AP-2 family of transcription factors. Genome Biol 6, 246.

Erickson, P. A., Baek, J., Hart, J. C., Cleves, P. A. and Miller, C. T. (2018). Genetic Dissection of a Supergene Implicates Tfap2a in Craniofacial Evolution of Threespine Sticklebacks. Genetics 209, 591–605.

Fan, X., Masamsetti, V. P., Sun, J. Q., Engholm-Keller, K., Osteil, P., Studdert, J., Graham, M. E., Fossat, N. and Tam, P. P. (2021). TWIST1 and chromatin regulatory proteins interact to guide neural crest cell differentiation. Elife 10.

Feng, Z., Duren, Z., Xiong, Z., Wang, S., Liu, F., Wong, W. H. and Wang, Y. (2021). hReg-CNCC reconstructs a regulatory network in human cranial neural crest cells and annotates variants in a developmental context. Commun Biol 4, 442.

Fernandez Garcia, M., Moore, C. D., Schulz, K. N., Alberto, O., Donague, G., Harrison, M. M., Zhu, H. and Zaret, K. S. (2019). Structural Features of Transcription Factors Associating with Nucleosome Binding. Mol Cell 75, 921–932 e926.

Fogelgren, B., Kuroyama, M. C., McBratney-Owen, B., Spence, A. A., Malahn, L. E., Anawati, M. K., Cabatbat, C., Alarcon, V. B., Marikawa, Y. and Lozanoff, S. (2008). Misexpression of Six2 is associated with heritable frontonasal dysplasia and renal hypoplasia in 3H1 Br mice. Dev Dyn 237, 1767–1779.

Gao, T., Wright-Jin, E. C., Sengupta, R., Anderson, J. B. and Heuckeroth, R. O. (2021). Cell-autonomous retinoic acid receptor signaling has stage-specific effects on mouse enteric nervous system. JCI Insight 6.

Gu, R., Zhang, S., Saha, S. K., Ji, Y., Reynolds, K., McMahon, M., Sun, B., Islam, M., Trainor, P. A., Chen, Y., et al. (2022). Single-cell transcriptomic signatures and gene regulatory networks modulated by Wls in mammalian midline facial formation and clefts. Development 149.

Han, J., Ishii, M., Bringas, P., Jr., Maas, R. L., Maxson, R. E., Jr. and Chai, Y. (2007). Concerted action of Msx1 and Msx2 in regulating cranial neural crest cell differentiation during frontal bone development. Mech Dev 124, 729–745.

Hao, Y., Hao, S., Andersen-Nissen, E., Mauck, W. M., 3rd, Zheng, S., Butler, A., Lee, M. J., Wilk, A. J., Darby, C., Zager, M., et al. (2021). Integrated analysis of multimodal single-cell data. Cell 184, 3573–3587 e3529.

Hari, L., Miescher, I., Shakhova, O., Suter, U., Chin, L., Taketo, M., Richardson, W. D., Kessaris, N. and Sommer, L. (2012). Temporal control of neural crest lineage generation by Wnt/beta-catenin signaling. Development 139, 2107–2117.

Heinz, S., Benner, C., Spann, N., Bertolino, E., Lin, Y. C., Laslo, P., Cheng, J. X., Murre, C., Singh, H. and Glass, C. K. (2010). Simple combinations of lineage-determining transcription factors prime cis-regulatory elements required for macrophage and B cell identities. Mol Cell 38, 576–589.

Hendershot, T. J., Liu, H., Clouthier, D. E., Shepherd, I. T., Coppola, E., Studer, M., Firulli, A. B., Pittman, D. L. and Howard, M. J. (2008). Conditional deletion of Hand2 reveals critical functions in neurogenesis and cell type-specific gene expression for development of neural crest-derived noradrenergic sympathetic ganglion neurons. Dev Biol 319, 179–191.

Hooper, J. E., Jones, K. L., Smith, F. J., Williams, T. and Li, H. (2020). An Alternative Splicing Program for Mouse Craniofacial Development. Front Physiol 11, 1099.

Hovland, A. S., Bhattacharya, D., Azambuja, A. P., Pramio, D., Copeland, J., Rothstein, M. and Simoes-Costa, M. (2022). Pluripotency factors are repurposed to shape the epigenomic landscape of neural crest cells. Dev Cell 57, 2257–2272 e2255.

Hsu, C. W., Wong, L., Rasmussen, T. L., Kalaga, S., McElwee, M. L., Keith, L. C., Bohat, R., Seavitt, J. R., Beaudet, A. L. and Dickinson, M. E. (2016). Three-dimensional microCT imaging of mouse development from early post-implantation to early postnatal stages. Dev Biol 419, 229–236.

Hufnagel, R. B., Zimmerman, S. L., Krueger, L. A., Bender, P. L., Ahmed, Z. M. and Saal, H. M. (2016). A new frontonasal dysplasia syndrome associated with deletion of the SIX2 gene. Am J Med Genet A 170A, 487–491.

Ishii, M., Han, J., Yen, H. Y., Sucov, H. M., Chai, Y. and Maxson, R. E., Jr. (2005). Combined deficiencies of Msx1 and Msx2 cause impaired patterning and survival of the cranial neural crest. Development 132, 4937–4950.

Iyyanar, P. P. R., Wu, Z., Lan, Y., Hu, Y. C. and Jiang, R. (2022). Alx1 Deficient Mice Recapitulate Craniofacial Phenotype and Reveal Developmental Basis of ALX1-Related Frontonasal Dysplasia. Front Cell Dev Biol 10, 777887.

Jacques-Fricke, B. T., Roffers-Agarwal, J. and Gammill, L. S. (2012). DNA methyltransferase 3b is dispensable for mouse neural crest development. PLoS One 7, e47794.

Jeong, J., Mao, J., Tenzen, T., Kottmann, A. H. and McMahon, A. P. (2004). Hedgehog signaling in the neural crest cells regulates the patterning and growth of facial primordia. Genes Dev 18, 937–951.

Kennedy, A. E. and Dickinson, A. J. (2012). Median facial clefts in Xenopus laevis: roles of retinoic acid signaling and homeobox genes. Dev Biol 365, 229–240.

Kenny, C., Dilshat, R., Seberg, H. E., Van Otterloo, E., Bonde, G., Helverson, A., Franke, C. M., Steingrimsson, E. and Cornell, R. A. (2022). TFAP2 paralogs facilitate chromatin access for MITF at pigmentation and cell proliferation genes. PLoS Genet 18, e1010207.

Kessler, S., Minoux, M., Joshi, O., Ben Zouari, Y., Ducret, S., Ross, F., Vilain, N., Salvi, A., Wolff, J., Kohler, H., et al. (2023). A multiple super-enhancer region establishes inter-TAD interactions and controls Hoxa function in cranial neural crest. Nat Commun 14, 3242.

Khor, J. M. and Ettensohn, C. A. (2020). Transcription Factors of the Alx Family: Evolutionarily Conserved Regulators of Deuterostome Skeletogenesis. Front Genet 11, 569314.

Knight, R. D., Javidan, Y., Nelson, S., Zhang, T. and Schilling, T. (2004). Skeletal and pigment cell defects in the lockjaw mutant reveal multiple roles for zebrafish tfap2a in neural crest development. Dev Dyn 229, 87–98.

Knight, R. D., Javidan, Y., Zhang, T., Nelson, S. and Schilling, T. F. (2005). AP2-dependent signals from the ectoderm regulate craniofacial development in the zebrafish embryo. Development 132, 3127–3138.

Knight, R. D., Nair, S., Nelson, S. S., Afshar, A., Javidan, Y., Geisler, R., Rauch, G. J. and Schilling, T. F. (2003). lockjaw encodes a zebrafish tfap2a required for early neural crest development. Development 130, 5755–5768.

Kowalczyk, M. S., Tirosh, I., Heckl, D., Rao, T. N., Dixit, A., Haas, B. J., Schneider, R. K., Wagers, A. J., Ebert, B. L. and Regev, A. (2015). Single-cell RNA-seq reveals changes in cell cycle and differentiation programs upon aging of hematopoietic stem cells. Genome Res 25, 1860–1872.

Kuleshov, M. V., Jones, M. R., Rouillard, A. D., Fernandez, N. F., Duan, Q., Wang, Z., Koplev, S., Jenkins, S. L., Jagodnik, K. M., Lachmann, A., et al. (2016). Enrichr: a comprehensive gene set enrichment analysis web server 2016 update. Nucleic Acids Res 44, W90–97.

Lakhwani, S., Garcia-Sanz, P. and Vallejo, M. (2010). Alx3-deficient mice exhibit folic acid-resistant craniofacial midline and neural tube closure defects. Dev Biol 344, 869–880.

Lamichhaney, S., Berglund, J., Almen, M. S., Maqbool, K., Grabherr, M., Martinez-Barrio, A., Promerova, M., Rubin, C. J., Wang, C., Zamani, N., et al. (2015). Evolution of Darwin’s finches and their beaks revealed by genome sequencing. Nature 518, 371–375.

Laugsch, M., Bartusel, M., Rehimi, R., Alirzayeva, H., Karaolidou, A., Crispatzu, G., Zentis, P., Nikolic, M., Bleckwehl, T., Kolovos, P., et al. (2019). Modeling the Pathological Long-Range Regulatory Effects of Human Structural Variation with Patient-Specific hiPSCs. Cell Stem Cell 24, 736–752 e712.

Lawson, N. D. and Weinstein, B. M. (2002). In vivo imaging of embryonic vascular development using transgenic zebrafish. Dev Biol 248, 307–318.

Li, H. and Durbin, R. (2009). Fast and accurate short read alignment with Burrows-Wheeler transform. Bioinformatics 25, 1754–1760.

Li, H., Handsaker, B., Wysoker, A., Fennell, T., Ruan, J., Homer, N., Marth, G., Abecasis, G., Durbin, R. and Genome Project Data Processing, S. (2009). The Sequence Alignment/Map format and SAMtools. Bioinformatics 25, 2078-2079.

Li, H., Sheridan, R. and Williams, T. (2013). Analysis of TFAP2A mutations in Branchio-Oculo-Facial Syndrome indicates functional complexity within the AP-2alpha DNA-binding domain. Hum Mol Genet 22, 3195–3206.

Li, W. and Cornell, R. A. (2007). Redundant activities of Tfap2a and Tfap2c are required for neural crest induction and development of other non-neural ectoderm derivatives in zebrafish embryos. Dev Biol 304, 338–354.

Lim, K. C., Lakshmanan, G., Crawford, S. E., Gu, Y., Grosveld, F. and Engel, J. D. (2000). Gata3 loss leads to embryonic lethality due to noradrenaline deficiency of the sympathetic nervous system. Nat Genet 25, 209–212.

Liu, Z., Li, C., Xu, J., Lan, Y., Liu, H., Li, X., Maire, P., Wang, X. and Jiang, R. (2019). Crucial and Overlapping Roles of Six1 and Six2 in Craniofacial Development. J Dent Res 98, 572–579.

Lohnes, D., Mark, M., Mendelsohn, C., Dolle, P., Dierich, A., Gorry, P., Gansmuller, A. and Chambon, P. (1994). Function of the retinoic acid receptors (RARs) during development (I). Craniofacial and skeletal abnormalities in RAR double mutants. Development 120, 2723–2748.

Love, M. I., Huber, W. and Anders, S. (2014). Moderated estimation of fold change and dispersion for RNA-seq data with DESeq2. Genome Biol 15, 550.

Luscher, B., Mitchell, P. J., Williams, T. and Tjian, R. (1989). Regulation of transcription factor AP-2 by the morphogen retinoic acid and by second messengers. Genes Dev 3, 1507–1517.

Mai, C. T., Isenburg, J. L., Canfield, M. A., Meyer, R. E., Correa, A., Alverson, C. J., Lupo, P. J., Riehle-Colarusso, T., Cho, S. J., Aggarwal, D., et al. (2019). National population-based estimates for major birth defects, 2010-2014. Birth Defects Res 111, 1420–1435.

Martik, M. L. and Bronner, M. E. (2021). Riding the crest to get a head: neural crest evolution in vertebrates. Nat Rev Neurosci.

Martino, V. B., Sabljic, T., Deschamps, P., Green, R. M., Akula, M., Peacock, E., Ball, A., Williams, T. and West-Mays, J. A. (2016). Conditional deletion of AP-2beta in mouse cranial neural crest results in anterior segment dysgenesis and early-onset glaucoma. Dis Model Mech 9, 849–861.

Matsuoka, T., Ahlberg, P. E., Kessaris, N., Iannarelli, P., Dennehy, U., Richardson, W. D., McMahon, A. P. and Koentges, G. (2005). Neural crest origins of the neck and shoulder. Nature 436, 347–355.

McGonnell, I. M., Graham, A., Richardson, J., Fish, J. L., Depew, M. J., Dee, C. T., Holland, P. W. and Takahashi, T. (2011). Evolution of the Alx homeobox gene family: parallel retention and independent loss of the vertebrate Alx3 gene. Evol Dev 13, 343–351.

McLean, C. Y., Bristor, D., Hiller, M., Clarke, S. L., Schaar, B. T., Lowe, C. B., Wenger, A. M. and Bejerano, G. (2010). GREAT improves functional interpretation of cis-regulatory regions. Nat Biotechnol 28, 495–501.

Meulemans, D. and Bronner-Fraser, M. (2002). Amphioxus and lamprey AP-2 genes: implications for neural crest evolution and migration patterns. Development 129, 4953–4962.

Milunsky, J. M., Maher, T. A., Zhao, G., Roberts, A. E., Stalker, H. J., Zori, R. T., Burch, M. N., Clemens, M., Mulliken, J. B., Smith, R., et al. (2008). TFAP2A mutations result in branchio-oculo-facial syndrome. Am J Hum Genet 82, 1171–1177.

Minoux, M., Holwerda, S., Vitobello, A., Kitazawa, T., Kohler, H., Stadler, M. B. and Rijli, F. M. (2017). Gene bivalency at Polycomb domains regulates cranial neural crest positional identity. Science 355.

Mitchell, J. M., Sucharov, J., Pulvino, A. T., Brooks, E. P., Gillen, A. E. and Nichols, J. T. (2021). The alx3 gene shapes the zebrafish neurocranium by regulating frontonasal neural crest cell differentiation timing. Development 148.

Muzumdar, M. D., Tasic, B., Miyamichi, K., Li, L. and Luo, L. (2007). A global double-fluorescent Cre reporter mouse. Genesis 45, 593–605.

Naqvi, S., Hoskens, H., Wilke, F., Weinberg, S. M., Shaffer, J. R., Walsh, S., Shriver, M. D., Wysocka, J. and Claes, P. (2022). Decoding the Human Face: Progress and Challenges in Understanding the Genetics of Craniofacial Morphology. Annu Rev Genomics Hum Genet 23, 383–412.

Neuhauss, S. C., Solnica-Krezel, L., Schier, A. F., Zwartkruis, F., Stemple, D. L., Malicki, J., Abdelilah, S., Stainier, D. Y. and Driever, W. (1996). Mutations affecting craniofacial development in zebrafish. Development 123, 357–367.

Pertea, M., Kim, D., Pertea, G. M., Leek, J. T. and Salzberg, S. L. (2016). Transcript-level expression analysis of RNA-seq experiments with HISAT, StringTie and Ballgown. Nat Protoc 11, 1650–1667.

Pertea, M., Pertea, G. M., Antonescu, C. M., Chang, T. C., Mendell, J. T. and Salzberg, S. L. (2015). StringTie enables improved reconstruction of a transcriptome from RNA-seq reads. Nat Biotechnol 33, 290–295.

Pimentel, H., Bray, N. L., Puente, S., Melsted, P. and Pachter, L. (2017). Differential analysis of RNA-seq incorporating quantification uncertainty. Nat Methods 14, 687–690.

Pini, J., Kueper, J., Hu, Y. D., Kawasaki, K., Yeung, P., Tsimbal, C., Yoon, B., Carmichael, N., Maas, R. L., Cotney, J., et al. (2020). ALX1-related frontonasal dysplasia results from defective neural crest cell development and migration. EMBO Mol Med 12, e12013.

Prescott, S. L., Srinivasan, R., Marchetto, M. C., Grishina, I., Narvaiza, I., Selleri, L., Gage, F. H., Swigut, T. and Wysocka, J. (2015). Enhancer divergence and cis-regulatory evolution in the human and chimp neural crest. Cell 163, 68–83.

Qu, S., Tucker, S. C., Zhao, Q., deCrombrugghe, B. and Wisdom, R. (1999). Physical and genetic interactions between Alx4 and Cart1. Development 126, 359–369.

Rada-Iglesias, A., Bajpai, R., Prescott, S., Brugmann, S. A., Swigut, T. and Wysocka, J. (2012). Epigenomic annotation of enhancers predicts transcriptional regulators of human neural crest. Cell Stem Cell 11, 633–648.

Ramirez, F., Ryan, D. P., Gruning, B., Bhardwaj, V., Kilpert, F., Richter, A. S., Heyne, S., Dundar, F. and Manke, T. (2016). deepTools2: a next generation web server for deep-sequencing data analysis. Nucleic Acids Res 44, W160–165.

Robinson, J. T., Thorvaldsdottir, H., Winckler, W., Guttman, M., Lander, E. S., Getz, G. and Mesirov, J. P. (2011). Integrative genomics viewer. Nat Biotechnol 29, 24–26.

Rothstein, M. and Simoes-Costa, M. (2020). Heterodimerization of TFAP2 pioneer factors drives epigenomic remodeling during neural crest specification. Genome Res 30, 35–48.

Roybal, P. G., Wu, N. L., Sun, J., Ting, M. C., Schafer, C. A. and Maxson, R. E. (2010). Inactivation of Msx1 and Msx2 in neural crest reveals an unexpected role in suppressing heterotopic bone formation in the head. Dev Biol 343, 28–39.

Santagati, F. and Rijli, F. M. (2003). Cranial neural crest and the building of the vertebrate head. Nat Rev Neurosci 4, 806–818.

Satoda, M., Zhao, F., Diaz, G. A., Burn, J., Goodship, J., Davidson, H. R., Pierpont, M. E. and Gelb, B. D. (2000). Mutations in TFAP2B cause Char syndrome, a familial form of patent ductus arteriosus. Nat Genet 25, 42–46.

Scerbo, P. and Monsoro-Burq, A. H. (2020). The vertebrate-specific VENTX/NANOG gene empowers neural crest with ectomesenchyme potential. Sci Adv 6, eaaz1469.

Schilling, T. F., Piotrowski, T., Grandel, H., Brand, M., Heisenberg, C. P., Jiang, Y. J., Beuchle, D., Hammerschmidt, M., Kane, D. A., Mullins, M. C., et al. (1996). Jaw and branchial arch mutants in zebrafish I: branchial arches. Development 123, 329–344.

Schmidt, M., Huber, L., Majdazari, A., Schutz, G., Williams, T. and Rohrer, H. (2011). The transcription factors AP-2beta and AP-2alpha are required for survival of sympathetic progenitors and differentiated sympathetic neurons. Dev Biol 355, 89–100.

Schorle, H., Meier, P., Buchert, M., Jaenisch, R. and Mitchell, P. J. (1996). Transcription factor AP-2 essential for cranial closure and craniofacial development. Nature 381, 235–238.

Seberg, H. E., Van Otterloo, E., Loftus, S. K., Liu, H., Bonde, G., Sompallae, R., Gildea, D. E., Santana, J. F., Manak, J. R., Pavan, W. J., et al. (2017). TFAP2 paralogs regulate melanocyte differentiation in parallel with MITF. PLoS Genet 13, e1006636.

Sekiguchi, R. and Hauser, B. (2019). Preparation of Cells from Embryonic Organs for Single-Cell RNA Sequencing. Curr Protoc Cell Biol 83, e86.

Selleri, L. and Rijli, F. M. (2023). Shaping faces: genetic and epigenetic control of craniofacial morphogenesis. Nat Rev Genet.

Simoes-Costa, M. and Bronner, M. E. (2016). Reprogramming of avian neural crest axial identity and cell fate. Science 352, 1570–1573.

Soldatov, R., Kaucka, M., Kastriti, M. E., Petersen, J., Chontorotzea, T., Englmaier, L., Akkuratova, N., Yang, Y., Haring, M., Dyachuk, V., et al. (2019). Spatiotemporal structure of cell fate decisions in murine neural crest. Science 364.

Soriano, P. (1999). Generalized lacZ expression with the ROSA26 Cre reporter strain. Nat Genet 21, 70–71.

Square, T., Jandzik, D., Romasek, M., Cerny, R. and Medeiros, D. M. (2017). The origin and diversification of the developmental mechanisms that pattern the vertebrate head skeleton. Dev Biol 427, 219–229.

Stenzel, A., Mumme-Monheit, A., Sucharov, J., Walker, M., Mitchell, J. M., Appel, B. and Nichols, J. T. (2022). Distinct and redundant roles for zebrafish her genes during mineralization and craniofacial patterning. Front Endocrinol (Lausanne) 13, 1033843.

Sucharov, J., Ray, K., Brooks, E. P. and Nichols, J. T. (2019). Selective breeding modifies mef2ca mutant incomplete penetrance by tuning the opposing Notch pathway. PLoS Genet 15, e1008507.

Tessier, P. (1976). Anatomical classification facial, cranio-facial and latero-facial clefts. J Maxillofac Surg 4, 69–92.

Tolarova, M. M. and Cervenka, J. (1998). Classification and birth prevalence of orofacial clefts. Am J Med Genet 75, 126–137.

van Dijk, D., Sharma, R., Nainys, J., Yim, K., Kathail, P., Carr, A. J., Burdziak, C., Moon, K. R., Chaffer, C. L., Pattabiraman, D., et al. (2018). Recovering Gene Interactions from Single-Cell Data Using Data Diffusion. Cell 174, 716–729 e727.

Van Otterloo, E., Feng, W., Jones, K. L., Hynes, N. E., Clouthier, D. E., Niswander, L. and Williams, T. (2016). MEMO1 drives cranial endochondral ossification and palatogenesis. Dev Biol 415, 278–295.

Van Otterloo, E., Li, H., Jones, K. L. and Williams, T. (2018). AP-2α and AP-2β cooperatively orchestrate homeobox gene expression during branchial arch patterning. Development 145.

Van Otterloo, E., Li, W., Bonde, G., Day, K. M., Hsu, M. Y. and Cornell, R. A. (2010). Differentiation of zebrafish melanophores depends on transcription factors AP2 alpha and AP2 epsilon. PLoS Genet 6, e1001122.

Van Otterloo, E., Li, W., Garnett, A., Cattell, M., Medeiros, D. M. and Cornell, R. A. (2012). Novel Tfap2-mediated control of soxE expression facilitated the evolutionary emergence of the neural crest. Development 139, 720–730.

Van Otterloo, E., Milanda, I., Pike, H., Thompson, J. A., Li, H., Jones, K. L. and Williams, T. (2022). AP-2alpha and AP-2beta cooperatively function in the craniofacial surface ectoderm to regulate chromatin and gene expression dynamics during facial development. Elife 11.

Vargel, I., Canter, H. I., Kucukguven, A., Aydin, A. and Ozgur, F. (2021). ALX-Related Frontonasal Dysplasias: Clinical Characteristics and Surgical Management. Cleft Palate Craniofac J, 10556656211019621.

Walker, M. B. and Kimmel, C. B. (2007). A two-color acid-free cartilage and bone stain for zebrafish larvae. Biotech Histochem 82, 23–28.

Wang, W. D., Melville, D. B., Montero-Balaguer, M., Hatzopoulos, A. K. and Knapik, E. W. (2011). Tfap2a and Foxd3 regulate early steps in the development of the neural crest progenitor population. Dev Biol 360, 173–185.

Wickham, H. (2016). ggplot2 : Elegant Graphics for Data Analysis. In Use R!,, pp. 1 online resource (XVI, 260 pages 232 illustrations, 140 illustrations in color. Cham: Springer International Publishing : Imprint: Springer,.

Williams, A. L. and Bohnsack, B. L. (2019). What’s retinoic acid got to do with it? Retinoic acid regulation of the neural crest in craniofacial and ocular development. Genesis 57, e23308.

Williams, T., Admon, A., Luscher, B. and Tjian, R. (1988). Cloning and expression of AP-2, a cell-type-specific transcription factor that activates inducible enhancer elements. Genes Dev 2, 1557–1569.

Williams, T. and Tjian, R. (1991). Characterization of a dimerization motif in AP-2 and its function in heterologous DNA-binding proteins. Science 251, 1067–1071.

Woodruff, E. D., Gutierrez, G. C., Van Otterloo, E., Williams, T. and Cohn, M. J. (2021). Anomalous incisor morphology indicates tissue-specific roles for Tfap2a and Tfap2b in tooth development. Dev Biol 472, 67–74.

Wu, Y., Kurosaka, H., Wang, Q., Inubushi, T., Nakatsugawa, K., Kikuchi, M., Ohara, H., Tsujimoto, T., Natsuyama, S., Shida, Y., et al. (2022). Retinoic Acid Deficiency Underlies the Etiology of Midfacial Defects. J Dent Res 101, 686–694.

Xie, Z., Bailey, A., Kuleshov, M. V., Clarke, D. J. B., Evangelista, J. E., Jenkins, S. L., Lachmann, A., Wojciechowicz, M. L., Kropiwnicki, E., Jagodnik, K. M., et al. (2021). Gene Set Knowledge Discovery with Enrichr. Curr Protoc 1, e90.

Xiong, Z., Dankova, G., Howe, L. J., Lee, M. K., Hysi, P. G., de Jong, M. A., Zhu, G., Adhikari, K., Li, D., Li, Y., et al. (2019). Novel genetic loci affecting facial shape variation in humans. Elife 8.

Xu, J., Liu, H., Lan, Y. and Jiang, R. (2022). The transcription factors Foxf1 and Foxf2 integrate the SHH, HGF and TGFbeta signaling pathways to drive tongue organogenesis. Development 149.

Xu, P., Balczerski, B., Ciozda, A., Louie, K., Oralova, V., Huysseune, A. and Crump, J. G. (2018). Fox proteins are modular competency factors for facial cartilage and tooth specification. Development 145.

Xu, P., Yu, H. V., Tseng, K. C., Flath, M., Fabian, P., Segil, N. and Crump, J. G. (2021). Foxc1 establishes enhancer accessibility for craniofacial cartilage differentiation. Elife 10.

Yoon, B., Yeung, P., Santistevan, N., Bluhm, L. E., Kawasaki, K., Kueper, J., Dubielzig, R., VanOudenhove, J., Cotney, J., Liao, E. C., et al. (2022). Zebrafish models of alx-linked frontonasal dysplasia reveal a role for Alx1 and Alx3 in the anterior segment and vasculature of the developing eye. Biol Open 11.

York, J. R. and McCauley, D. W. (2020). The origin and evolution of vertebrate neural crest cells. Open Biol 10, 190285.

Zalc, A., Rattenbach, R., Aurade, F., Cadot, B. and Relaix, F. (2015). Pax3 and Pax7 play essential safeguard functions against environmental stress-induced birth defects. Dev Cell 33, 56–66.

Zhang, J., Hagopian-Donaldson, S., Serbedzija, G., Elsemore, J., Plehn-Dujowich, D., McMahon, A. P., Flavell, R. A. and Williams, T. (1996). Neural tube, skeletal and body wall defects in mice lacking transcription factor AP-2. Nature 381, 238–241.

Zhao, Q., Behringer, R. R. and de Crombrugghe, B. (1996). Prenatal folic acid treatment suppresses acrania and meroanencephaly in mice mutant for the Cart1 homeobox gene. Nat Genet 13, 275–283.

